# MethylCal: Bayesian calibration of methylation levels

**DOI:** 10.1101/587519

**Authors:** Eguzkine Ochoa, Verena Zuber, Nora Fernandez-Jimenez, Jose Ramon Bilbao, Graeme R. Clark, Eamonn R. Maher, Leonardo Bottolo

## Abstract

Bisulfite amplicon sequencing has become the primary choice for single-base methylation quantification of multiple targets in parallel. The main limitation of this technology is a preferential amplification of an allele and strand in the PCR due to methylation state. This effect, known as “PCR bias”, causes inaccurate estimation of the methylation levels and calibration methods based on standard controls have been proposed to correct for it. Here, we present a Bayesian calibration tool, MethylCal, which can analyse jointly all CpGs within a DMR or CpG island, avoiding “one-at-a-time” CpG calibration. This enables more precise modeling of the methylation levels observed in the standard controls. It also provides accurate predictions of the methylation levels not considered in the controlled experiment, a feature that is paramount in the derivation of the corrected methylation degree. We tested the proposed method on eight independent assays (two CpG islands and six imprinting DMRs) and demonstrated its benefits, including the ability to detect outliers. We also evaluated MethylCal’s calibration in two practical cases, a clinical diagnostic test on 18 patients potentially affected by Beckwith-Wiedemann syndrome, and 17 individuals with celiac disease. The calibration of the methylation levels obtained by MethylCal allows a clearer identification of patients undergoing loss or gain of methylation in borderline cases and could influence further clinical or treatment decisions. MethylCal is availability as an R package on https://github.com/lb664/MethylCal.

## Introduction

DNA methylation is an epigenetic mark associated with a broad range of disorders including cancer (1), autoimmunity (2), aging (3) and imprinting (4). This mechanism implies the addition of a methyl group to the 5’-carbon of cytosine in a CpG dinucleotide to form 5-methylcytosine (5-mC) (5). Modifications in DNA methylation could affect gene expression as reported in several types of diseases (6–9).

To validate epigenome associations, identify region of interest or clinically relevant biomarkers and create new diagnostic tests, it is crucial to develop fast, cheap and accurate DNA methylation assays (10). In this sense, bisulfite amplicon sequencing is an ideal choice for its capacity to analyse multiple targets in parallel with high accuracy, concordance and low cost (11). However, this method critically requires the amplification of bisulfite converted DNA for the discrimination between un-methylated and methylated cytosines. The bisulfite conversion consists in the modification of un-methylated cytosines on uracil (U) maintaining methylated cytosines as cytosines (C). The result of the conversion is a single strand fragmented DNA no longer complemented. If there is a preferential amplification of an allele and strand in the PCR, this effect is called “PCR bias” (12). In order to obtain accurate results, it is important to minimize its effect as much as possible. To this end, investigators (13, 14) have proposed to redesign primers by looking at strand-specific as well as bisulfite-specific flanking primers, but this solution is expensive and time consuming and might not solve the problem completely. Instead, PCR bias can be calculated and corrected *in silico* (12, 15) by using standard controls with known methylation levels. Specifically, the best-fit hyperbolic (12) and cubic polynomial (15) curve obtained from the apparent level of methylation after PCR in standard controls is used to correct the observed methylation levels in the case and control samples.

In this work, we propose a new Bayesian calibration method that overcomes the limitations of the existing tools. In particular, our method analyses jointly all CpGs within a CpG island (CGI) or a Differentially Methylated Region (DMR), avoiding “one-at-a-time” CpG calibration or the calibration of the average methylation level across CpGs that neglects the variability across CpGs (15, 16). To test the proposed method, we designed eight independent assays in two CGIs located on *SDHC* gene promoter and six imprinted DMRs, see Table 1 for details. After genomic DNA bisulfite conversion, each target region was amplified by specific primers, and specific amplicons were sequenced on MiSeq. Each assay was run on five standard controls with known methylation percentages (0%, 25%, 50%, 75% and 100%) to determine the specific calibration curve through MethylCal. Compared to existing calibration tools (12,15), our method is able to capture with precision the variability of the apparent level of methylation observed after amplification at different actual methylation percentages. We demonstrate this feature and the benefits of our method when deriving the calibration curves in all the assays analysed.

**Table 1.**
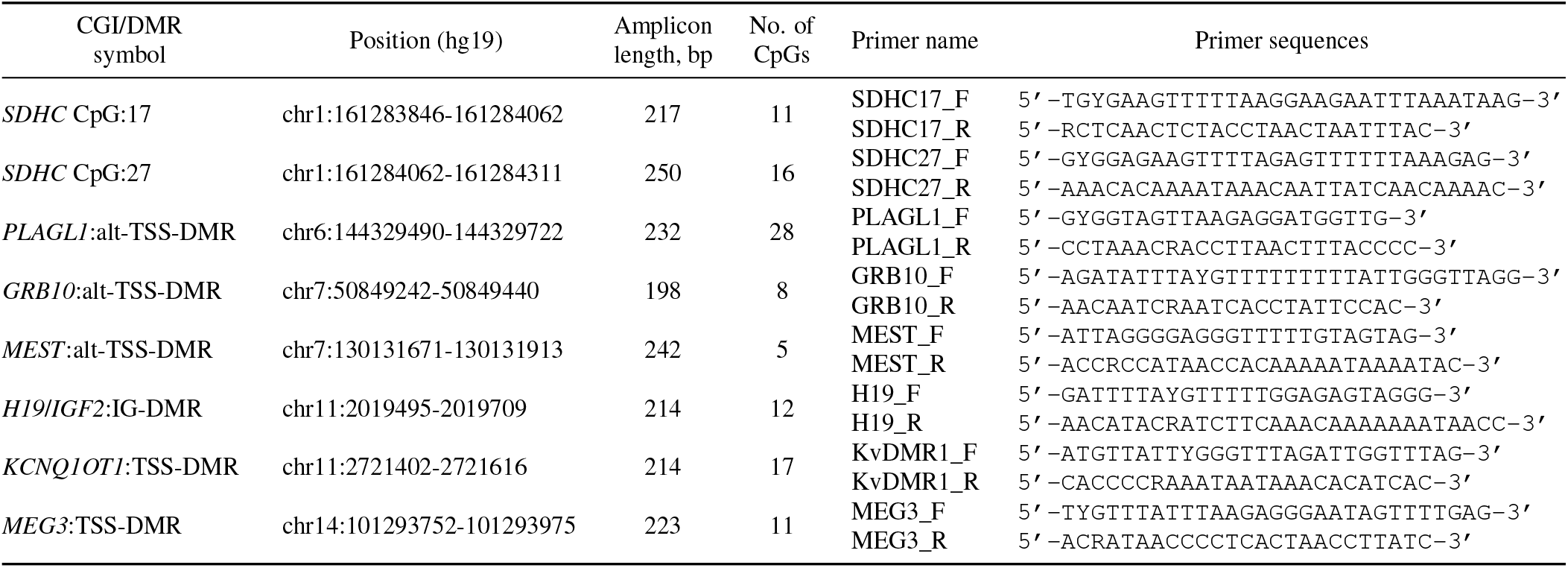
Sequences of the PCR primers used in this study (TSS for transcription start site, IG for intergenic, and alt-TSS for alternative transcription start site). For easy of notation, in the main text, tables and figures’ caption, and supplementary material, we refer only to the gene name instead of the CGI/DMR symbol.

When applied to a data set consisting of 18 patients potentially affected by Beckwith-Wiedemann syndrome (BWS) (17), the calibration curves obtained by our new method permit a more precise correction of the observed levels of methylation in two target regions (*KCNQ10T1* and *H19/IGF2*) with a clearer identification of patients undergoing loss or gain of methylation. We also validated MethylCal in a second data set regarding patients with celiac disease (16,18). Our method achieved better calibrations and more reliable corrections of the methylation levels in three target regions that have been associated with susceptibility to celiac disease. These features are important in clinical practice, since the accurate calibration of the methylation levels obtained by more sophisticated statistical methods could influence treatment decisions or further actions.

## MATERIALS AND METHODS

### Samples

For standard controls, we used Human methylated and non-methylated DNA set from Zymo (Zymo, CA, USA). The non-methylated DNA was purified from HCT116 DKO cells knockout for both DNA methyltransferases DNMT1 (−/−) and DNMT3b (−/−). The methylated DNA was purified from the same HCT116 DKO cells and was enzymatically methylated by M.SssI methyltransferase. Five actual methylation percentage (0%, 25%, 50%, 75% and 100%) were prepared mixing different ratios of non-methylated and methylated human control DNA (Zymo, CA, USA) bisulphite converted (MethylEdge Bisulfite Conversion System, Promega). Additionally, we collected DNA from 18 potential BWS patients and 15 healthy controls. Genomic DNA was extracted from peripheral blood using Gentra Puregene Blood Kit (Qiagen) and DNA quality was determined by Qubit 2.0 (Invitrogen, ThermoFisher). Appropriate human subject approvals and written inform consent were obtained from all participants. Bisulphite conversion of genomic DNA was performed in all samples at the same time with with MethylEdge Bisulfite Conversion System from Promega.

### PCR amplification

We designed eight assays to quantify the methylation level at each CpG site in two CGIs located on *SDHC* gene promoter and six imprinted DMRs, see Table 1 for details. For the design of the primers we used Bisulfite Primer Seeker 12S a tool developed by Zymo (http://bpsbackup.zymoresearch.com/). The primers parameters were: 20-32 bp primer length, 150-220 bp product length, 55-57°C Tm, allowing 1 CpG in the first 1/3 of primer, whereas the minimum number of CpGs per product is 4. All designs were tested for primer dimers by the Multiple Primer Analyzer software (ThermoFisher). To allow sequencing through Nextera XT kit (Illumina), we added overhangs sequences to each primer, forward overhang 5’–TCGTCGGCAGCGTCAGATGTGTATAAGAGACAG–3’ and reverse overhang 5’–GTCTCGTGGGCTCGGAGATGTG TATAAGAGACAG–3’. All standard controls were bisulfite converted at the same time and eight specific PCRs were run at the same time using the same standard controls. To determine the conversion rate, we examined the conversion of cytosines on thymidine in non-CpG sites in non-methylated control DNA (0%) showing a conversion rate higher than 98% (19, 20). The sequences of the PCR primers used are listed in Table 1. The PCR reactions were carried out in 25 μl with ZymoTaq Premix (Zymo) using 1.2 μl of bisulphite converted DNA. The amplification program was 95°C for 10 min, then 40 cycles at 95°C for 30 sec, 58°C for 40 seg and 72°C for 1 min, and elongation step at 72°C for 7 min. PCR products were purified with QIAquick PCR Purification Kit (Qiagen). To attached dual indices and Illumina sequencing adapters we performed a second PCR of the purified products using Nextera XT index kit (Illumina) following the recommendations of manufacturer’s. The second PCR was purified with AMPure XP beads (Beckman Coulter), quantified by Qubit 2.0 (Invitrogen) and normalized to 4 nM.

### Bisulphite sequencing

Fastq files obtained by MiSeq system (Illumina) were trimmed with *cutadapt* software (http://cutadapt.readthedocs.io/en/stable/) using quality (`–q 30`) and short reads (`–m 50`) parameters. Trimmed reads were aligned with *bismark* software using human reference build hg19 (https://www.bioinformatics.babraham.ac.uk/projects/bismark/). Finally, to extract the methylation level per every single cytosine, we used the tool *bismark_methylation_extractor* included in the *bismark* package.

### MethylCal

#### Model outline

MethylCal is a fully Bayesian mixed additive regression model. It predicts the apparent level of methylation observed after amplification based on the actual methylation percentages (AMP), borrowing information across all CpGs within a CGI or a DMR. MethylCal’s regression model can be described as follows

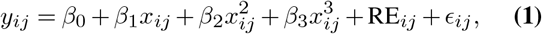

where *y_ij_* ∈ [0%, 100%] is the apparent level of methylation after PCR at the *i*th AMP (*i* =1,…,*l*) and the *j*th CpG (*j* = 1,…,*m*), *x_ij_* is the ith AMP (*x*_1*j*_ = 0% and *x_lj_* = 100%) which is constant across CpGs (in our experimental design 0%, 25%, 50%, 75% and 100% actual methylation are the same for all CpGs) and *β*_0_,…,*β*_3_ are the coefficients of the polynomial regression. Finally, *ϵ_ij_* ~ *N*(0,*σ*^2^).

MethylCal is based on Moskalev’s cubic polynomial regression (CPR) (15) given its simplicity, flexibility and effectiveness to calibrate methylation data. However, instead of fitting a distinct CPR for each CpG, in (1) CpGs are jointly analysed using all *n* = *l × m* observations at once. The second key feature of our model is the inclusion of the random-effects RE_*ij*_(*i* =1,…,*l, j* = 1,…,*m*) that capture distinct effects at each AMP or CpG or a combination of both. Depending on how RE_*ij*_ is defined, different models can be derived from (1). In the next section, we present the specification of RE_*ij*_ that we found useful in order to model accurately the apparent level of methylation after PCR in standard controls.

#### Random-effects specification

MethylCal includes four regression models that differ by the specification of the random-effects RE_*ij*_ and the model that fits better the data is selected by the Deviance Information Criterion (DIC) (21). The regression models considered in MethylCal are

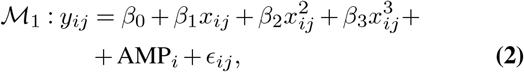

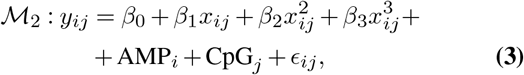

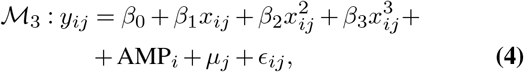

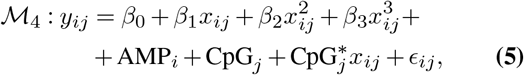

Besides the fixed-effects polynomial regression terms, in (2) the random-effects AMP_*i*_ are introduced to model the variability of the apparent level of methylation after PCR at different AMPs not explained by the CPR. In (3) the crossed random-effects (22) CpG_*j*_ are added to capture the heterogeneity of the apparent level of methylation across CpGs. In (4) the latent Gaussian field (LGF) ***μ*** = (*μ*,…,*μ_m_*)^*T*^ (23) replaces the crossed random-effects CpG_*j*_ to model the dependence of the apparent levels of methylation across CpGs. Finally, in (5) the random-slopes (22) 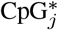 are added to model the larger (smaller) variability of the apparent level of methylation after PCR across CpGs at lower (higher) AMPs. The opposite scenario with a smaller (larger) variability at lower (higher) AMPs is also considered in (5). For identifiability conditions, in all models considered, we assume that ∑_*i*_AMP_*i*_ = 0, ∑_*j*_CpG_*j*_ = 0 and 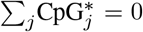. Supplementary Figure S.1 provides a schematic representation of MethylCal’s regression models, highlighting the role of the CPR, the crossed random-effects AMP_*i*_ and CpG_*j*_, the LGF ***μ*** and the combined effect of the random-intercepts CpG and random-slopes CpG* in predicting the apparent level of methylation after PCR.

#### Priors set-up

Since MethylCal is a fully Bayesian model, a prior distribution is specified for each unknown parameter. The fixed-effects regression coefficients follow a non-informative normal prior distribution, *β*_1_,…,*β*_4_ ~ N(0,10^3^), whereas for the intercept an improper prior distribution is used, 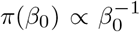. The crossed random-effects AMP_*i*_ and CpG_*j*_ follow a normal distribution, AMP_*i*_|*τ* ~ N(0,*τ*^−1^) and CpG_*j*_|*υ* ~ N(0,*υ*^−1^) with a non-informative prior precision *τ* ~ Gam(1,0.1) (E(*τ*) = 10 and Var(*τ*) = 100) and *υ* ~ Gam(1,0.1). For the LGF, we follow (24) and model ***μ*** as a Random Walk of order 1 (RW1) (25), *μ_j_*|*μ*_*j*−1_,*ρ_j_* ~ N(*μ*_*j*−1_,*ρ_j_*), where *ρ_j_* = *ρ*|*p_j_* − *p*_*j*−1_| with *ρ*^−1^ ~ Gam(1,0.1) and *p_j_* and *p*_*j*−1_ the chromosomal position of two consecutive CpGs (with *p*_0_ = 0). With this specification the dependence between methylation levels depends on the distance between the corresponding CpGs, i.e., the closer the CpGs, the stronger the dependence. Finally, for the random-intercepts/slopes model, we specify a Normal-Wishard prior, 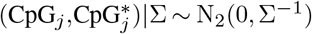, with Σ ~ W_2_(*r,R*) (E(Σ^−1^) = *R*/(*r* − 3)). The default values for the Wishart hyperparameters are *r* = 4 and *R* = *I*_2_. The prior set-up is completed by the specification of a proper but relatively uninformative prior on the error variance, *σ*^2^ ~ InvGam(10^−10^,0.001).

#### Advantages of the proposed model

MethylCal has several advantages compared to existing calibration tools. First, CpGs are jointly analysed using all *n* = *l × m* observations at once, avoiding unrealistic assumptions of independence of the methylation levels at nearby CpGs. Second, MethylCal is more parsimonious with fewer parameters to estimate (five for the main effects, including the error variance, and *l* + 1, *l* + *m* + 2, and *l* + 2*m* + 4 random-effects coefficients for model 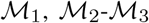 and 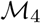, respectively, in contrast to 5*m* coefficients required by Moskalev’s CPR, where *m* is the number of CpGs in a DMR or CGI). Combined with a larger sample size, it allows narrower coefficients credible intervals and smaller prediction credible intervals, i.e., less model uncertainty. Third, differently from a simple fixed-effects model, the specification of different random effects allows MethylCal to adequately account for the patterns of variances and correlations of the methylation levels. While in Moskalev’s method 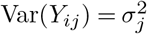 is constant across AMPs, MethylCal allows a more complex variance structure. In model 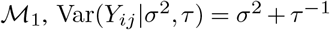, where *τ*^−1^ models the variability of the apparent level of methylation after PCR across AMPs. In model 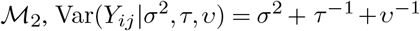, Var(*Y_ij_*|*σ*^2^,*τ, υ*) = *σ*^2^ + *τ*^−1^ + *υ*^−1^ with *υ*^−1^ the additional variability of the apparent level of methylation across CpGs, whereas in model 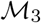, the RW1 induces the autoregressive variance decomposition 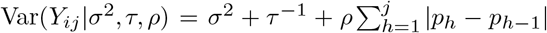. In model 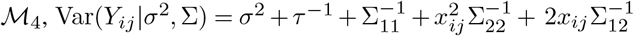 with 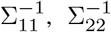 and 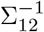 the elements of the covariance matrix Σ^−1^. Finally, in contrast to Moskalev’ CPR, MethylCal is able to capture the dependence between the observations and, in particular, the dependence of the methylation levels across CpGs (26). For instance in model 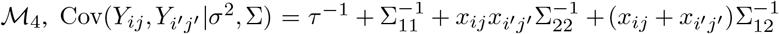 with *Y_ij_* and *Y*_*i*′*j*′_ the observations from two distinct AMPs and CpGs and with *x_ij_* and *x*_*i*′*j*′_ the corresponding actual methylation percentages.

#### Inference

Inference on MethylCal’s parameters is performed using INLA R package (http://www.r-inla.org/). INLA is a probabilistic language that performs approximate Bayesian inference by means of integrated nested Laplace approximations (27) and numerical integrations. The main advantage of INLA is its simplicity since a known practical impediment of Monte Carlo Markov chain methods in real applications is the large computational burden. Instead, INLA only requires the specification of the regression model, similarly to other regression packages in R (https://www.r-project.org/). A second advantage is its computational speed since no sampling is required from the posterior densities. This is particularly important in model 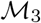 since LGF posterior inference is rather difficult using Monte Carlo Markov chain.

Let *β* = (*β*_0_,*β*_1_,*β*_2_,*β*_3_), *γ* = (*σ*^2^,*τ,υ*, Σ) and ***μ*** be the vector of the fixed effects, the vector of variance components and the LGF, respectively. Integrating out the random-effects AMP_*i*_, CpG_*j*_ and 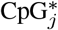, (2)–(5) can be rewritten in a more compact formulation that encompasses all models considered

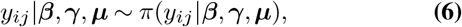

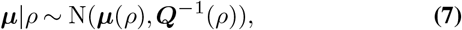

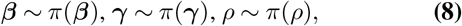

where (6) is the observations equation, (7) is the latent Gaussian field equation with mean ***μ***(*ρ*) and sparse precision matrix ***Q***(*ρ*) and (8) are the parameters equations. INLA inferential procedure for MethylCal’s models consists of three steps:

1. Compute the approximation to the marginal posterior *π*(***β***,*γ,ρ*|***y***) and by-product to *π*(***β|y***), *π*(*γ*|***y***) and *π*(*ρ*|***y***);
2. Compute the approximation to *π*(*μ_j_*|***y***,*ρ*);
3. Combine 1. and 2. above to compute 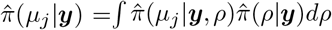, where 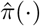 is the approximated density.

Note that steps 2 and 3 are only required for model 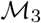. Despite the Laplace approximations and numerical integrations, INLA provides results that are very close to those obtained by exact MCMC methods. Details about INLA procedure can be found in (28) and (23).

Given the additive structure of MethylCal, the predictive values are derived straightforwardly. For example, in model 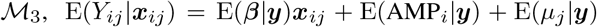, E(*Y_ij_*|***x***_*ij*_) = E(***β|y***)***x***_*ij*_ +E(AMP_*i*_|***y***) + E(*μ_j_*|***y***), E(*Y_ij_*|***x***_*ij*_) ∈ [0%, 100%], with 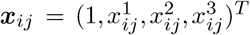, E(***β|y***) the posterior mean of the fixed effects, E(AMP_*i*_|*y*) the posterior mean of the random-effects AMP at the *i*th level and E(*μ_j_*|***y***) the posterior mean of the LGF at the *j*th CpG. Similarly, *q_α_*(*Y_ij_*|***x***_*ij*_) = *q_α_*(***β|y***)***x***_*ij*_ + *q_α_*(AMP_*i*_|***y***) + *q_α_*(*μ_j_*|***y***), where *q_α_*(·) ∈ [0%, 100%] is the *α*% quantile of the posterior distribution.

### Predictive measures

We compare the predictive ability of the MethylCal’s model selected by the DIC with Moskalev’s CPR. In particular, we report the following “in-sample” and “out-of-sample” predictive measures:

- Residual Sum of Squares: RSS = ∑_*ij*_{*y_ij_* − E(*Y_ij_*|***x***)}^2^;
- Mean Squared Error of Prediction (29): MSEP = Σ_*ij*_{*y_ij_* − E(*Y_ij_*|***x***_\(*ij*)_)}^2^/*n*, where E(*Y_ij_*|***x***_\(*ij*)_) indicates the prediction of *y_ij_* when the observation corresponding to the *i*th AMP *and j*th CpG is excluded from the regression. We also consider the case E(*Y_ij_*|***x***_*i\j*_) when the *j* th CpG is removed and E(*Y_ij_*|***x***_\*i,j*_) when the *i*th AMP is excluded from all CpGs;
- CV-index (30): 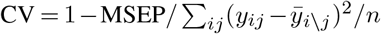, where 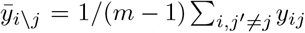 is the average apparent level of methylation after PCR without the measurement corresponding to the *j*th CpG. MSEP is the Mean Squared Error of Prediction when ***x***_\(*ij*)_ or ***x***_*i\j*_ are removed from the regression. The case ***x***_\*i,j*_ is not considered.

The RSS ∈ [0,1] is a measure of “in-sample” fit and it is well known that over-parameterized models achieve usually better RSS. The MSEP ∈ [0, +∞) is instead a measure of “out-of-sample” prediction based on leave-one-out cross-validation. A model with lower MSEP should be preferred since it predicts more accurately the apparent level of methylation after PCR for unobserved values of the actual methylation percentages, a feature that is important in the derivation of the corrected methylation degree. Finally, the CV-index ∈ (−∞, 1] is similar to MSEP, but it aims at comparing the “out-of-sample” prediction of the proposed model with a simpler non-parametric model that predicts the apparent level of methylation after PCR by using the average value of all other observations. A negative CV-index is in favour of a simpler non-parametric model versus a more sophisticated parametric one.

### Corrected methylation degree

Given an observed level of methylation, measured in an individual (either in the case or in the control group) at a particular CpG within DMR or a CGI, the corrected methylation degree can be obtained. However, differently from (12), where it can be calculated analytically by inverting the equation that describes the calibration curve, both Moskalev’s CPR and MethylCal require a numerical procedure to perform the PCR-bias correction. In (15), the corrected methylation degree is obtained by solving

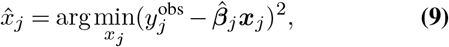

where 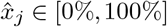 is the corrected methylation degree, 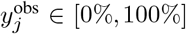 is the observed level of methylation at the *j*th CpG, 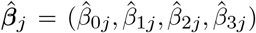 is the maximum likelihood solution of the CPR for the *j*th CpG based on apparent level of methylation after PCR in the standard controls and 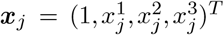. The existence of an unique solution depends on 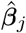, but in the examples considered Moskalev’s CPR is a strict increasing function. Thus, the objective function (9) admits only one solution which can be obtained by the R function *optimize*.

The derivation of the objective function for MethylCal’s mixed additive regression model is slightly more complicated since only few known values of the actual methylation percentage are usually tested in a calibration experiment. This is a typical problem in linear mixed models when the predictions are made for new observations, as these predictions are conditional on an unobserved level of the random effect (31). To overcome this problem, we consider *η_i_* = E(AMP_*i*_|***y***) the posterior mean of the random-effects AMP at the ith level. Note that *η_i_* is also the predicted value of the random-effects AMP at the same ith level of the observation *x_ij_*. A cubic spline interpolation is then fitted on the posterior means *η_i_*(*i* = 1,…,*l*) and, by doing so, a new value *η*(*x_j_*) can be predicted for any value of the actual methylation percentage *x_j_* ∈ [0%, 100%], see Supplementary Figure S.8 for illustrative examples of the cubic spline interpolation of the posterior mean of the random-effects AMP. Finally, for each CpG (*j* = 1,…,*m*), the PCR-bias corrected methylation degree 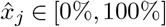 is the solution of

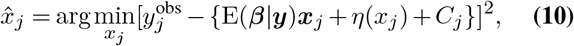

where E(***β|y***) is the posterior mean of the fixed effects, *η*(*x_j_*) is the cubic spline predicted value of the random-effects AMP at the new observation *x_j_, C_j_* = 0, *C_j_* = E(CpG_*j*_|***y***), *C_j_* = E(*μ_j_*|***y***) and 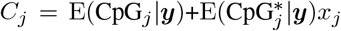 for model 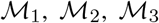 and 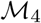, respectively. In (10) the existence of an unique solution depends on the combined effect of E(***β|y***), *η*(*x_j_*) and *C_j_*, but in the examples analysed, MethylCal’s calibration curve is strictly increasing, allowing for an unique solution. For each CpG, the numerical value of 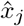 is then obtained by using the R function *optimize*, specifying (10) as the objective function.

## RESULTS

The assays presented in Table 1 were analysed using MethylCal and Moskale’s CPR. First, for each specific assay, we derived the calibration curves using five standard controls with known methylation percentages. Secondly, we checked the goodness of fit of the calibration curves obtained by MethylCal and compared the results with those obtained by Moskalev’s CPR. Thirdly, we corrected the observed methylation degree based on the estimated calibration curves in two specific target regions (*KCNQ1OT1* and *H19/IGF2*) important for their BWS clinical diagnostic value and in three target genes that have been associated with susceptibility to celiac disease. The three steps are detailed below.

### Derivation of the calibration curves

We obtained specific calibration curve for each assay using the proposed method and compared the results with Moskalev’s CPR. Figure 1 shows the level of methylation of two assays *KCNQ1OT1* (top panels) and *H19/IGF2* (bottom panels) predicted by Moskalev’s CPR, whereas Figure 2 shows the results obtained by MethylCal. Moskalev’s CPR is an over-parameterized model: when the actual methylation percentages (AMP) at 0%, 25%, 50%, 75% and 100% are used in the calibration experiment, the number of estimated parameters for each CpG (four for the regression coefficients and one for the residual variance) equals the number of observations, leaving no degrees of freedom. Thus, the 95% prediction confidence interval is extremely wide, see Figures 1B-E, with a large uncertainty regarding the estimated model. In contrast, Figures 2B-E highlight the parsimony of MethylCal. Using all *n* = *l × m* observations at once, with less parameters to estimate and thus higher degrees of freedom, the 95% prediction credible interval are much smaller than Moskalev’s CPR. Moreover, using MethylCal, the predicted level of methylation are very close to the apparent level of methylation after PCR. This is evident by looking at Figures 2A-D, where model 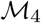 and 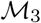 were selected by the DIC for the assay *KCNQ1OT1* and *H19/IGF2*. In both assays, MethylCal interpolates the apparent level of methylation observed after amplification remarkably well despite the fact that the data show a more complex pattern than previously reported hyperbolic (12) or cubic polynomial (15) shape when the apparent level of methylation after PCR is plotted as a function of the actual methylation percentage. In contrast, in Figures 1A-D, Moskalev’s CPR is not able to interpolate the data with the same precision, in particular for the inner values of the AMPs (25%, 50%, 75%). Finally, Figures 1C-F and Figures 2C-F show the impact of the interpolation on the PCR-bias correction. For each CpG-AMP combination, (9) and (10) are used to correct the apparent level of methylation after PCR. If the correction is perfect, the corrected methylation degrees will coincide with the AMPs used in the calibration experiment. Overall, MethylCal’s correction is more precise than Moskalev’s adjustment due to its ability to interpolate adequately the apparent level of methylation after PCR at different AMPs.

**Figure 1.**
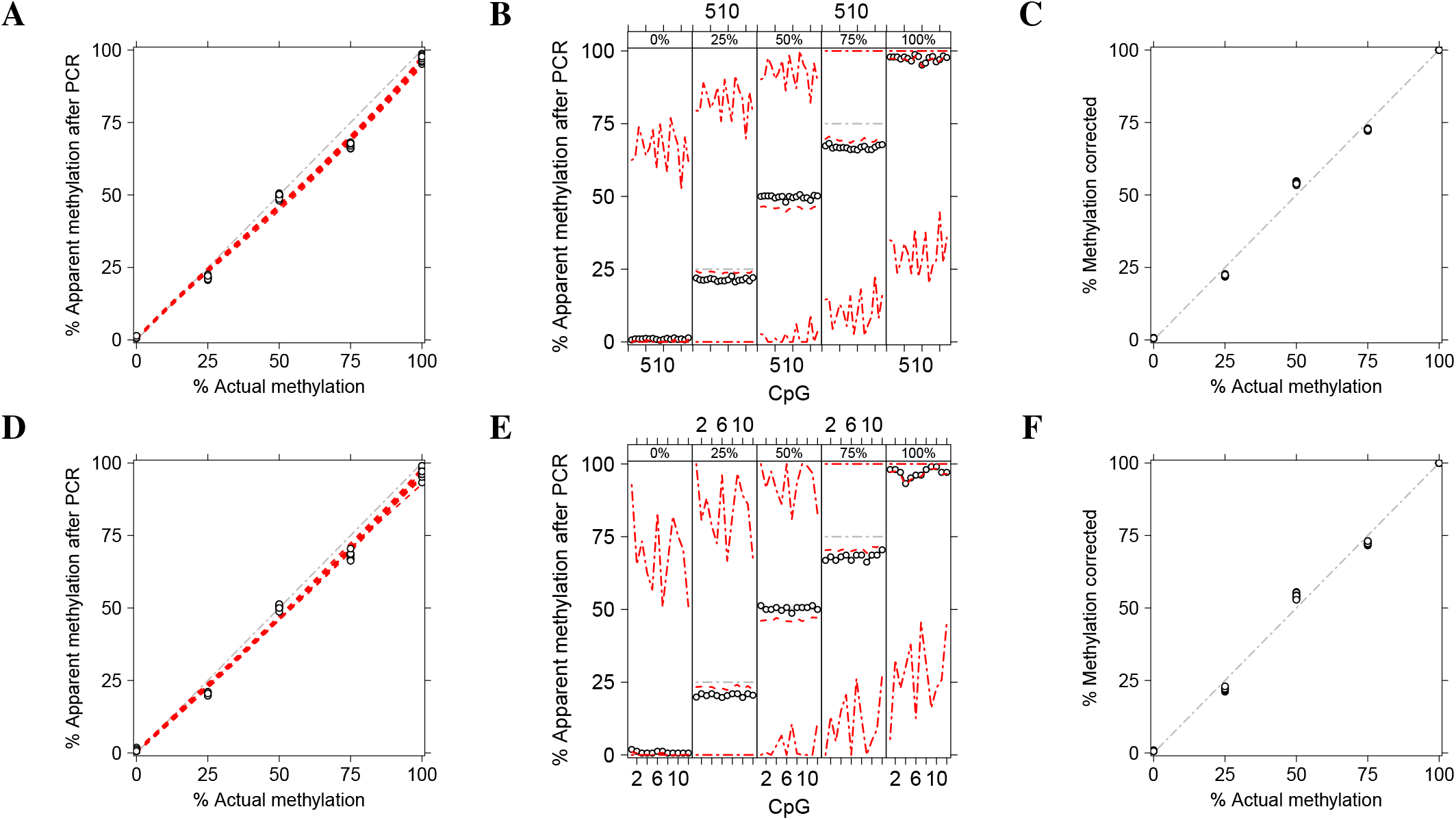
Methylation level of two independent assays *KCNQ1OT1* (top panels) and *H19/IGF2* (bottom panels) calibrated by Moskalev’s cubic polynomial regression (CPR). (**A-D**) The apparent level of methylation observed after amplification (*y*-axis) is plotted as a function of the actual methylation percentage (AMP) (*x*-axis). Circles depict the apparent level of methylation after PCR for each CpG at different AMPs, whereas the red dotted lines show Moskalev’s CPR for each CpG. The grey dash-dotted line represents an unbiased plot. (**B-E**) The apparent level of methylation observed after amplification (*y*-axis) is plotted (circles) as a function of the CpGs in the DMR (*x*-axis), stratified by AMP (top figure box and the grey dot-dashed line). For each stratum, the red dotted line shows the level of methylation predicted by Moskalev’s CPR, whereas the red dot-dashed lines depict the 95% prediction confidence interval. (**C-F**) The result of PCR-bias correction by using Moskalev’s CPR. The corrected methylation degree (*y*-axis) is plotted as a function of the AMP (*x*-axis). The grey dash-dotted line represents an unbiased corrected plot.

**Figure 2.**
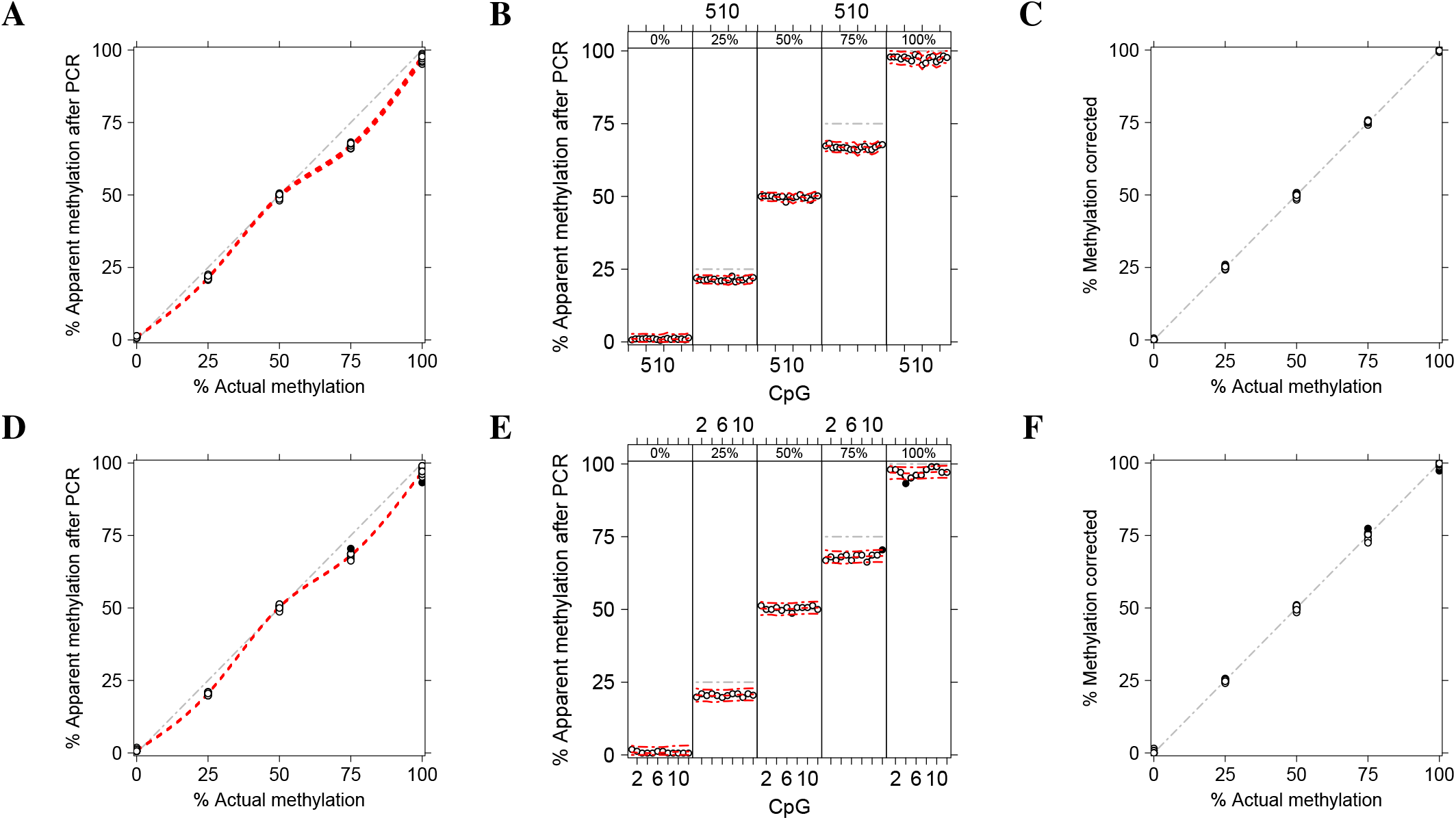
Methylation level of two independent assays *KCNQ1OT1* (top panels) and *H19/IGF2* (bottom panels) calibrated by MethylCal with model 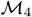 and 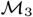 selected by the DIC, respectively. (**A-D**) The apparent level of methylation observed after amplification (*y*-axis) is plotted as a function of the actual methylation percentage (AMP) (*x*-axis). Circles depict the apparent level of methylation after PCR for each CpG at different AMPs, whereas the red dotted lines show MethylCal’s predicted level of methylation for each CpG. Black dots highlight potential outliers. The grey dash-dotted line represents an unbiased plot. (**B-E**) The apparent level of methylation observed after amplification (*y*-axis) is plotted (circles) as a function of the CpGs in the DMR (*x*-axis), stratified by AMP (top figure box and the grey dot-dashed line). For each stratum, the red dotted line shows the level of methylation predicted by MethylCal, whereas the red dot-dashed lines depict the 95% prediction credible interval. (**C-F**) The result of PCR-bias correction obtained by using MethylCal. The corrected methylation degree (*y*-axis) is plotted as a function of the AMP (*x*-axis). The grey dash-dotted line represents a unbiased corrected plot.

By visual inspection of MethylCal’s results presented in Figure 2F, some measurements seem less well calibrated at 75% actual methylation. A closer look at Figure 2E reveals that the apparent level of methylation after PCR for CpG 12 is outside the posterior predictive interval [*l,u*] for outliers detection with *l* = Q1 − 1.5IQR and *u* = Q3 + 1.5IQR with IQR = Q3 − Q1 and Q3 and Q1 the 75th and 25th percentiles of the posterior predictive density. A second CpG outside the posterior predictive interval for outliers detection is present also at 100% actual methylation although in this case, given the shape of the PCR-bias correction curve, the impact on the calibration is less pronounced. Under the fitted MethylCal’s model, these observations can be either regarded as outliers, and thus removed from the analysis, or the data generation process, including biological and biochemical factors, should be further investigated to understand the possible causes of this unusual pattern. This conclusion highlights a further feature of MethylCal, i.e., its ability to pinpoint specific CpG-AMP combinations as potential outliers that do not fit with the bulk of the data and need to be further checked.

The predicted level of methylation and the PCR-bias correction for the rest of the assays analysed, two CGIs (*SDHC* CpG:17 and *SDHC* CpG:27) located on SDHC gene promoter and four imprinted DMRs (*PLAGL1, GRB10, MEST* and *MEG3*) are shown in Supplementary Figures S.4 and S.5 for Moskalev’s CPR and in Supplementary Figures S.6 and S.7 for MethylCal. The results are similar to those presented in Figures 1 and 2 with larger prediction credible intervals, less accurate predicted level of methylation and worse corrected methylation degree when Moskalev’s CPR is used. These conclusions are similar across different assays, including the previously reported cubic polynomial data shape as shown for the *PLAGL1* assay in Supplementary Figures S.4G and S.6G.

Finally, by visual inspection of the data, three CpGs (21-23) were removed from the *PLAGL1* assay since their pattern was extremely different from other observations in the same assay. While Moskalev’s CPR cannot impute missing CpGs, MethylCal is able to impute them, see Supplementary Figure S.6H, based on the fixed-effects terms (which are common to all CpGs) and the posterior mean of the random-effects AMP, see Supplementary Figure S.8C.

### Goodness of fit

MethylCal’s superior performance compared to Moskalev’s CPR is also apparent when the “in-sample” and “out-of-sample” goodness-of-fit measures are considered. Table 2 shows the predictive performance for two assays tested, *KCNQ1OT1* and *H19/IGF2*. The best MethylCal’s model selected by the DIC performs better than Moskalev’s CPR when the Residual Sum of Squares (RSS) is considered. These results demonstrate that, although Moskalev’s CPR is an over-parameterised model that should attain better “in-sample” prediction, it is not suitable for calibration when the data do not show previously reported hyperbolic or cubic polynomial data shapes. The same conclusions can be drawn for the other assays presented in Supplementary Table S.1. MethylCal shows better predictive performance in all assays tested and only marginally worst for the *GRB10* assay. MethylCal performs better also in the *PLAGL1* assay which is the most favourable case for Moskalev’s CPR given the cubic polynomial shape of the data.

**Table 2.**
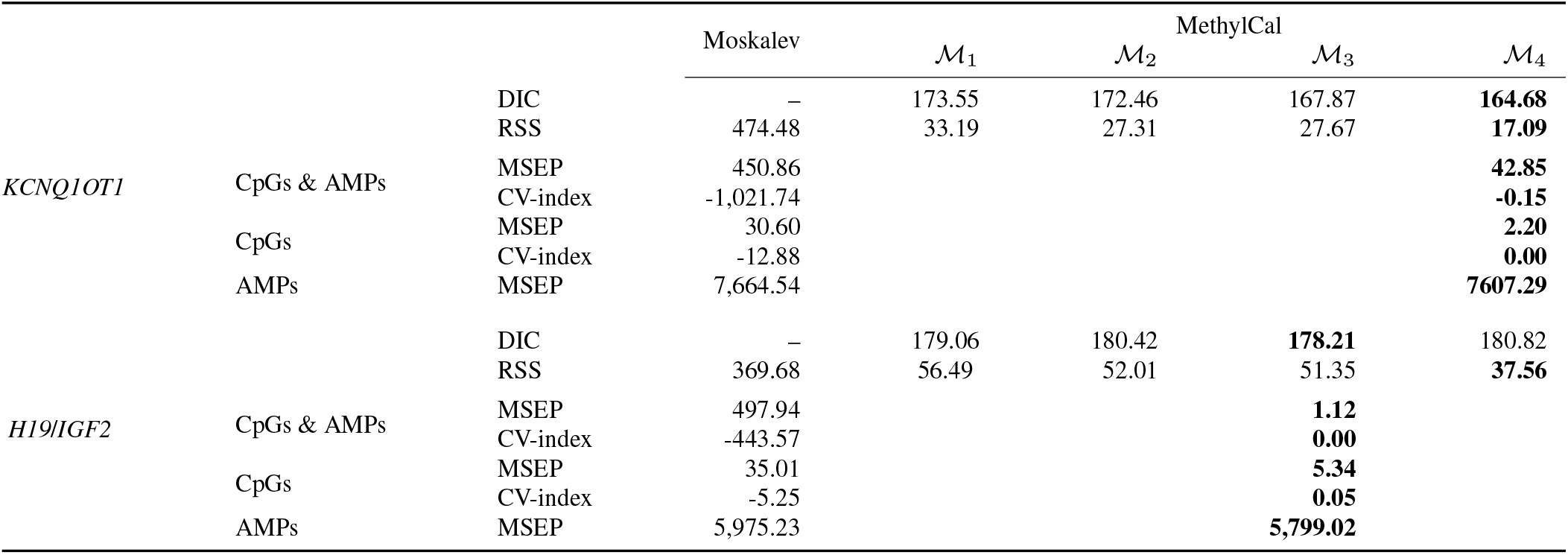
Goodness-of-fit performance of MethylCal and Moskalev’s cubic polynomial regression (CPR) on two independent assays *KCNQ1OT1* and *H19*/*IGF2*. For each MethylCal’s model the Deviance Information Criterion (DIC) is calculated and for MethylCal’s models and Moskalev’s CPR also the Residual Sum of Squares (RSS). The Mean Squared Error of Prediction (MSEP) and the CV-index are evaluated for the best MethylCal’s model (based on the DIC) and for Moskalev’s CPR when the leave-one-out cross-validation is performed across (i) CpGs and actual methylation percentages (AMPs); (ii) CpGs; (iii) AMPs. For each predictive measure, the best result is highlighted in bold.

Our comparisons also consider the “out-of-sample” prediction and three possible scenarios are examined. In the first one, the cross-validation is performed by removing a data point that corresponds to a specific CpG-AMP combination. In the second scenario, each CpG is excluded one-at-a-time, while in the last scenario each AMP is removed separately in the cross-validation. Since Moskalev’s method cannot predict the AMPs of the CpGs that have been removed, in the second scenario the “out-of-sample” prediction is obtained by averaging Moskalev’s predicted values of the two flanking CpGs, each of them weighted by the distance (in bp) between the excluded CpG and each flanking CpG.

In all assays tested, the Mean Squared Error of Prediction (MSEP) of the best identified MethylCal’s model is lower than Moskalev’s CPR by several orders of magnitude when the cross-validation is performed across CpG-AMP combinations or across CpGs, see Table 2 and Supplementary Table S.1. When the cross-validation is performed across AMPs the difference between MethylCal and Moskalev’s CPR is less evident. Since in this scenario an AMP has been removed from all CpGs, MethylCal cannot borrow information about the excluded AMP across CpGs. Nonetheless, MethylCal has lower MSEP than Moskalev’s method across all assays analysed, with a gain ranging between 1% and 5%. The improvement for the *GRB10* assay (around 43%) is particularly high since in this case the exclusion of a calibration sample does not hurt the estimation of the random-effects AMP_*i*_, see Supplementary Figure S.8E. Taken together, these results suggest that MethylCal should be also preferred when the number of calibration samples is reduced from five to four.

We also evaluate MethylCal’s performance by using the CV-index. Interestingly, the CV-index for Moskalev’s CPR is always negative when a CpG-AMP combination is removed in the cross-validation. Thus, a non-parametric model that predicts the CpG-AMP combination by using the remaining observations performs better than Moskalev’s CPR. This is also true when a CpG is excluded in the cross-validation, but the *GRB10* assay. In contrast, when looking at the CV-index, selected the best MethylCal’s model is always better than Moskalev’s CPR and it has an inferior CV-index performance only in one case (*KCNQ1OT1* assay) in the prediction of the CpG-AMP combination and another one (*SDHC CpG:17*) in the CpG “out-of-sample” prediction.

Finally, Supplementary Table S.3 summarises MethylCal’s goodness-of-fit measures across all assays tested and compares them with Moskalev’s CPR. MethylCal’s best model selected by the DIC performs always better than Moskalev’s CPR in the “in-sample” prediction, but in a single assay. In the “out-of-sample” prediction MethylCal’s best model is always better than Moskalev’s CPR (with the exception of the *GRB10* assay) either considering the MSEP or the CV-index measures. Moreover, MethylCal’s best model has a non-negative CV-index in 14 out of 16 cases.

### Application in clinical diagnostic of Beckwith-Wiedemann syndrome

BWS is caused by genetic and epigenetic abnormalities on chr11p15.5-–11p15.4 that produce an increment of IGF2 growth factor levels and/or a reduction of CDKN1C growth suppressor protein levels. The loss of methylation of maternal *KCNQ1OT1* and the gain of methylation of maternal *H19*/*IGF2* are the most frequent defects in BWS. In addition, the frequency of mosaicism is high in BWS, introducing the problem of borderline cases that are difficult to diagnose.

The observed methylation levels of 15 healthy controls and 18 potential BWS patients were corrected using the calibration curves obtained by MethylCal and Moskalev’s CPR in the *KCNQ1OT1* and *H19*/*IGF2* assays. Patients with an average corrected methylation level below a 3SD confidence interval were considered to undergo loss of methylation and those with a level above the 3SD confidence interval were considered to experience gain of methylation, see Figures 3A-B and 4A-B for the assays *KCNQ1OT1* and *H19*/*IGF2*, respectively, using Moskalev’s CPR (left panels) and MethylCal (right panels). To avoid false positives, in clinical practice a ±3SD confidence interval is usually chosen since it guarantees low type-I error (*α* =0.0027). Moreover, the confidence interval should be large enough to contain the control samples’ corrected methylation degrees across all CpGs.

**Figure 3.**
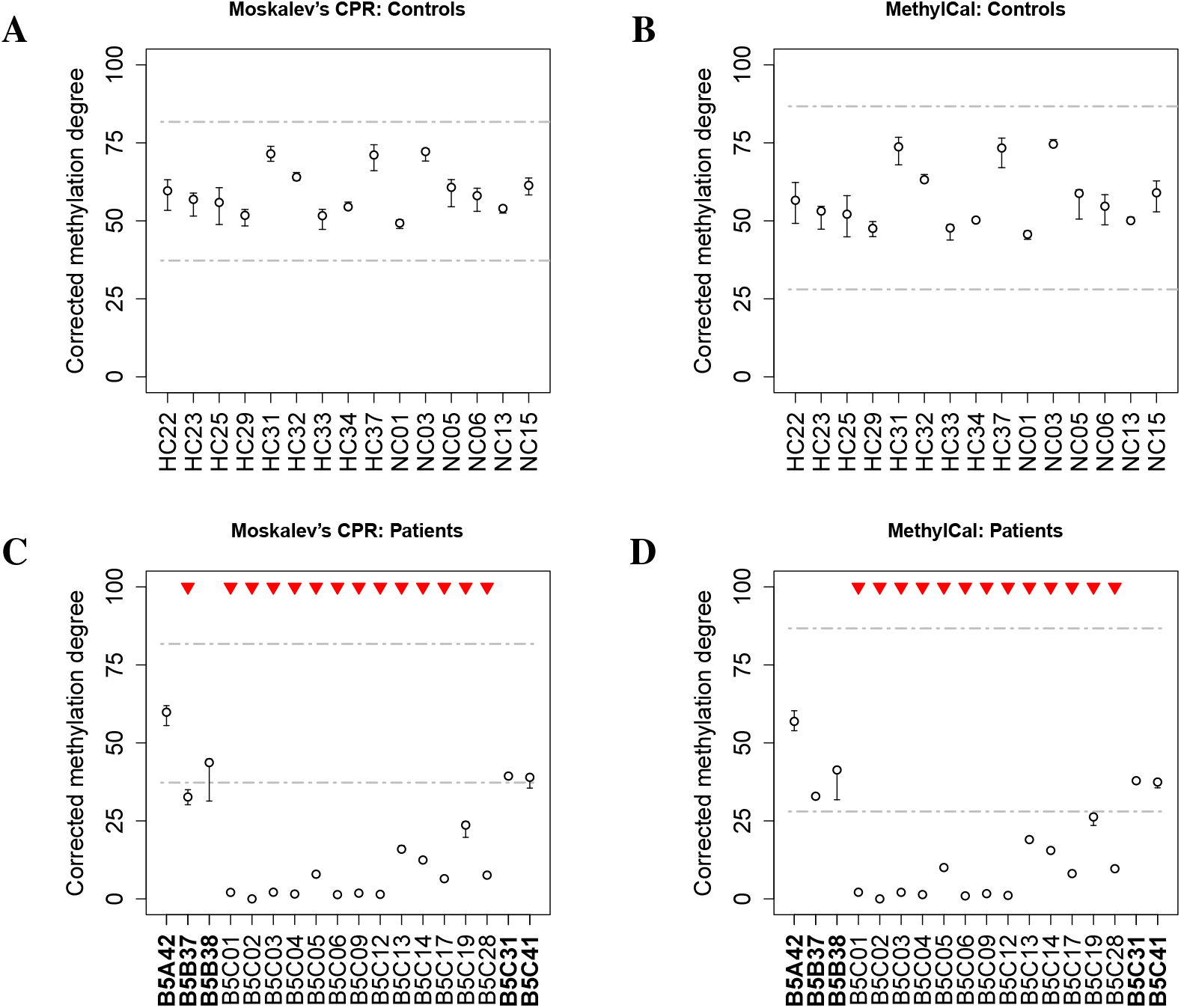
Corrected methylation degree of the *KCNQ1OT1* assay using Moskalev’s cubic polynomial regression (left panels) and MethylCal (right panels) for healthy controls (top panels) and patients potentially affected by Beckwith-Wiedemann syndrome (bottom panels). (**A-B**) For each healthy control (*x*-axis), the boxplot depicts the range and the median of the corrected methylation degree (*y*-axis) across CpGs. The dashed-dotted grey lines show the ±3SD confidence interval centered around the overall mean (see main text for details). (**C-D**) For each patient (*x*-axis), the boxplot depicts the range and the mean (circle) of the corrected methylation degree (*y*-axis) across CpGs while the dashed-dotted grey lines show the healthy controls’ confidence interval. Top red triangles indicate patients classified as having undergone loss or gain of methylation if their average (across CpGs) corrected methylation degree is outside the healthy controls’ confidence interval. Bold font (*x*-axis) indicates patients’ classification described in the main text.

**Figure 4.**
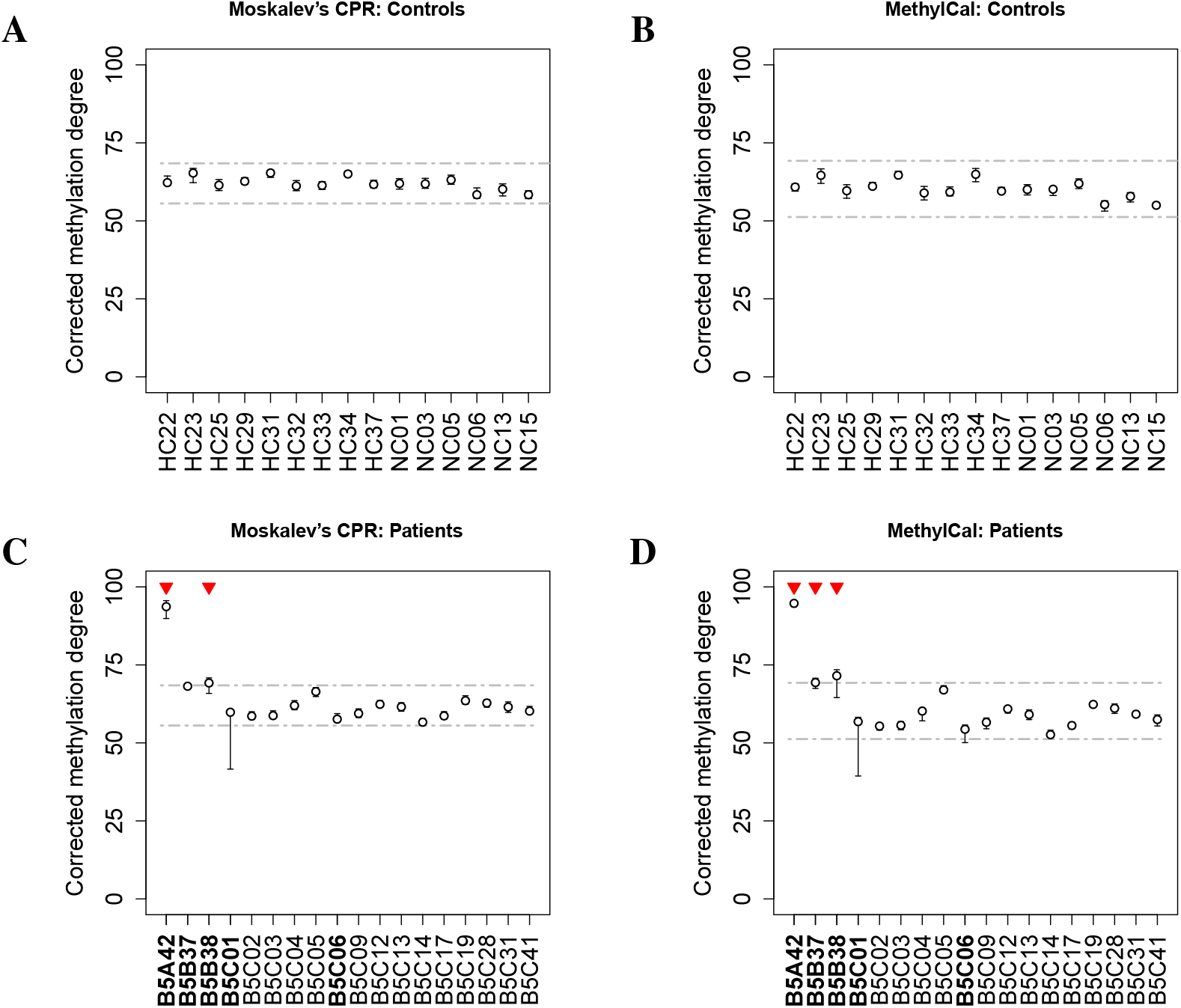
Corrected methylation degree of the *H19/IGF2* assay using Moskalev’s cubic polynomial regression (left panels) and MethylCal (right panels) for healthy controls (top panels) and patients potentially affected by Beckwith-Wiedemann syndrome (bottom panels). (**A-B**) For each healthy control (*x*-axis), the boxplot depicts the range and the median of the corrected methylation degree (*y*-axis) across CpGs. The dashed-dotted grey lines show the ±3SD confidence interval centered around the overall mean (see main text for details). (**C-D**) For each patient (*x*-axis), the boxplot depicts the range and the mean (circle) of the corrected methylation degree (*y*-axis) across CpGs while the dashed-dotted grey lines show the healthy controls’ confidence interval. Top red triangles indicate patients classified as having undergone loss or gain of methylation if their average (across CpGs) corrected methylation degree is outside the healthy controls’ confidence interval. Bold font (*x*-axis) indicates patients’ classification described in the main text.

Figures 3A-B present the results of the corrected methylation degree for the *KCNQ1OT1* assay in the healthy control group. MethylCal has a larger confidence interval (28.012-86.71) compared to that obtained by using Moskalev’s CPR (37.26-81.73). This is due to the effect of the calibration curve estimated by Moskalev’s CPR that shrinks the corrected methylation degrees for observed methylation levels greater than 50%, while the opposite happens for observed methylation levels lower than 50%, see Figure 1C. The joint effect of a larger healthy controls’ confidence interval and a more accurate calibration of the methylation degree in the patients group permit to reclassify patients B5B37 as normal methylated in contrast to Moskalev’s CPR that classifies the same patient as having undergone loss of methylation, see Figures 3C-D. Moreover, with MethylCal, patients B5B38 and B5C41 are well within the healthy controls’ confidence interval (including the range of the corrected methylation degree across CpGs) with less uncertainty about their classification.

Figures 4C-D show the results of a second assay, *H19*/*IGF2*, used in the classification of patients. While both methods detected gain of methylation in patients B5A42 and B5B38, and thus affected by BWS, patient B5B37 is also identified as having undergone gain of methylation by MethylCal. However, in contrast to the *KCNQ1OT1* assay, in the *H19*/*IGF2* assay there is more uncertainty regarding the classification: for both patients B5B37 and B5B38, the range of the corrected methylation degree across CpGs intersects the upper bound of the healthy controls’ confidence interval, while normal-classified methylated patients B5C01 and B5C06 show the same uncertainty at the bottom of the confidence interval.

Finally, patients’ classification depends upon the choice of the length of the healthy controls’ confidence interval. However, when a less conservative test is chosen (*α* = 0.01), MethylCal’s results do not change. This is not true when Moskalev’s CPR is employed as shown in Supplementary Figure S.11. This is due to Moskalev’s less precise calibration curve and its shrinkage effect on the corrected methylation degrees for which a small difference in the level of significance has a large impact on the patients’ classification.

### Application in clinical diagnostic of celiac patients

We applied MethylCal in a second data set containing human genomic control DNA measured at eight distinct AMPs (0%, 12.5%, 25%, 37.5%, 50%, 62.5%, 87.5% and 100%) in eight NFkB-related and Toll-like receptor genes (16). It also contains the uncorrected methylation levels on the same target regions of 13 controls and 17 celiac patients at the time of diagnosis with patient data pyrosequenced in three runs (18). In our analysis we focused on *NFKBIA* gene, as well as on *RELA* and *TNFAIP3* genes that, similarly to *NFKBIA*, have been associated with susceptibility to celiac disease.

Figure 5 shows the calibration curves of the *NFKBIA* assay and the corrected methylation degrees in celiac patients using Moskalev’s CPR (left panels) and MethylCal (right panels). Our method confirms its ability to automatically detect outliers. For example, in Figures 5 B-D, several methylation levels in CpG 5 are detected as outliers (black dots) since they show an apparent level of methylation at 37.5%, 50% and 62.5% actual methylation that is lower than at 25%. Similarly, there is an outlier in CpG 3 where the apparent level of methylation at 50% actual methylation is as high as at 62.5%. Outliers were also detected between 37.5% and 62.5% actual methylation in the *RELA* and *TNFAIP3* assays, see Figures S.12B-D and S.13B-D, respectively. Rather than relying on a difficult visual inspection of the data, MethylCal identifies specific CpG-AMP combinations that do not fit with the bulk of the data and it accounts for them when it derives the calibration curves. See also Supplementary Table S.3 for the comparison of the goodness of fit between MethylCal and Moskalev’s CPR on the *NFKBIA, RELA* and *TNFAIP3* assays and the overall better performance of the proposed tool.

**Figure 5.**
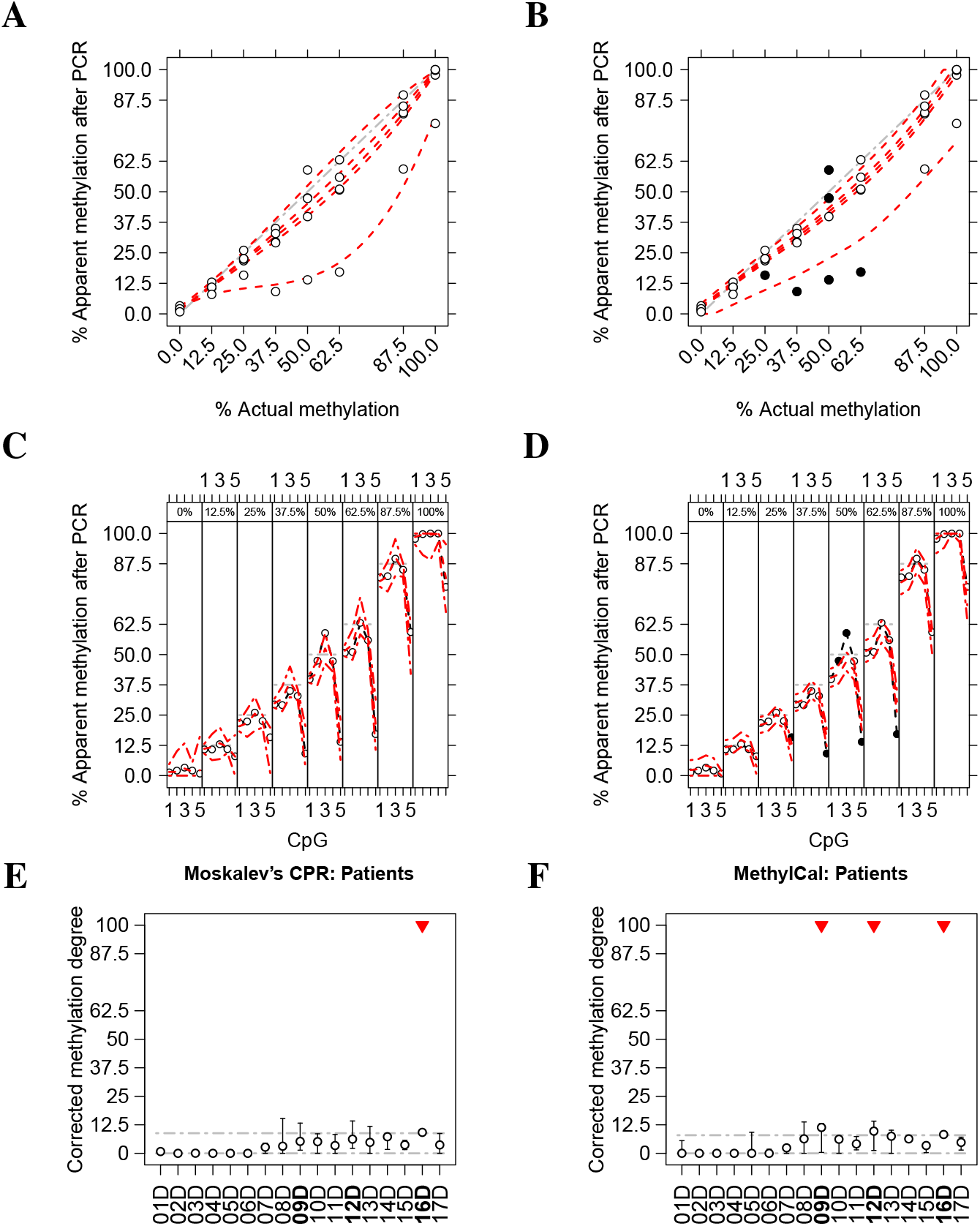
Calibrated methylation level and corrected methylation degree of the *NFKBIA* assay in celiac patients using Moskalev’s cubic polynomial regression (left panels) and MethylCal (right panels). (**A-B**) The apparent level of methylation observed after amplification (*y*-axis) is plotted as a function of the actual methylation percentage (AMP) (*x*-axis). Circles depict the apparent level of methylation after PCR for each CpG at different AMPs, whereas the red dotted lines show MethylCal’s predicted level of methylation for each CpG. Black dots highlight potential outliers. The grey dash-dotted line represents an unbiased plot. (**C-D**) The apparent level of methylation observed after amplification (*y*-axis) is plotted (circles) as a function of the CpGs in the DMR (*x*-axis), stratified by AMP (top figure box and the grey dot-dashed line). For each stratum, the red dotted line shows the level of methylation predicted by MethylCal, whereas the red dot-dashed lines depict the 95% prediction credible interval. (**E-F**) For each patient (*x*-axis), the boxplot depicts the range and the mean (circle) of the corrected methylation degree (*y*-axis) across CpGs while the dashed-dotted grey lines show the healthy controls’ confidence interval. Top red triangles indicate patients classified as having undergone loss or gain of methylation if their average (across CpGs) corrected methylation degree is outside the healthy controls’ confidence interval. Bold font (*x*-axis) indicates patients’ classification described in the main text.

A different estimation of the calibration curves may have a large impact on the correction of the case/controls samples and the classification of the patients. Figures 5E-F exemplify this case where Moskalev’s CPR classifies patient 16D as normal methylated, while MethylCal, besides patient 16D, identifies patients 09D and 12D as normal methylated. In particular, MethylCal estimates an average corrected methylation level for patient 09D (11.34) that is more than double the level obtained by Moskalev’s CPR (5.24). Further investigations confirm that patient 09D is always classified by Moskalev’s CPR as having undergone loss of methylation irrespectively of the level of significance of the test (*α* ≤ 0.10).

## DISCUSSION

Bisulfite amplicon sequencing is an ideal platform for the detection of methylation changes on multiple targets in parallel due to the low cost and the efficiency in the single-base quantification (32). The main limitation of this technology is a preferential amplification of an allele and strand in the PCR due to methylation state (12). This effect causes inaccurate estimation of the methylation and *in silico* calibration tools have been proposed to minimize it.

In this work, we proposed a new Bayesian calibration tool that is able to analyse jointly all CpGs within a CGI or a DMR avoiding “one-at-a-time” CpG calibration. MethylCal has several benefits compared to existing methods (12, 15), including a better “out-of-sample” prediction which is particularly important in the derivation of the corrected methylation degree and the ability to detect CpG-AMP combinations that should be regarded as outliers, and therefore removed or further checked. Our approach is also very general and it is applicable irrespectively of the locus analysed (CGIs or DMRs), the type and degree of PCR bias to be recovered (large towards the un-methylated allele as in the *PLAGL1* assay, small towards the methylated allele as in *SDHC* CpG:17), the number of CpGs per locus (few as in the *MEST* assay, many as in the *PLAGL1* assay) and the number of calibration samples.

MethylCal includes four different models, each of which with a different random-effects combination. In the analysis of BWS data, 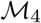 is the preferred model since it allows the specification of the correlation of the apparent level of methylation between CpGs and the AMPs. This behaviour is particularly evident in two assays, *GRB10* and *MEST*, that show higher than expected methylation levels at 0% and 25% actual methylation, an effect that gradually disappears at higher AMPs (Supplementary Figures S.2C-E). Although less pronounced, the opposite behaviour is present in the assays *H19/IGF2* and *KCNQ1OT1* (Figures S.2G-I) and *SDHC* CpG:27 (Supplementary Figure S.3B). This pattern cannot be explained by the expected error associated with standard controls (5% for un-methylated DNA and an extra 5% for fully-methylated DNA). Given that the same calibration samples were used in all reactions and the high conversion rate (> 98%), this phenomenon might be due to regions specific resistance to bisulfite conversion. A possible explanation is the formation of stable secondary structures around the CpG site that makes the region more resistant to denaturation and subsequent conversion (33).

The application of MethylCal for the calibration of the observed methylation levels of the *KCNQ1OT1* and *H19/IGF2* assays in a real data set of possible BWS cases shows the importance of the accurate quantification and correction of the PCR bias to distinguish borderline cases. We considered gain of methylation in a region when the level of methylation is above 3SD from the average methylation level detected in the control group and loss of methylation when the level of methylation is below 3SD. Using MethylCal, we classified patients B5B37 as having undergone gain of methylation in the target region *H19/IGF2* in contrast to Moskalev’s method that identified the same patient with a loss of methylation in the target region *KCNQ1OT1*. In the analysis, we applied a very conservative threshold, ±3SD (around 0.3% of false-positive error in the diagnostic), but MethylCal’s results do not change if the level of significance is increased to a less restrictive 1%, demonstrating that its corrections are less influenced by the choice of the level of the test. Finally, the benefits of the proposed method, i.e., better calibrations and more reliable corrections, are also shown in a second case/control data set related to pyrosequenced methylation levels in three target regions associated with susceptibility to celiac disease.

In both real data applications, the improvement in the accuracy observed after calibration determines the diagnosis, but it could also influence clinical or treatment decisions or further actions. Moreover, the accuracy of the calibration method is critical in disorders with mosaicism like BWS but not exclusive, since the same problem will affect, for example, circulating tumor DNA samples, which will have extensive application in cancer diagnostic in a near future.

In conclusion, MethylCal learns the existence, location and size of PCR bias better than existing methods and adjusts for it in the correction step, allowing the identification of loss or gain of methylation in difficult cases with less uncertainty compared to existing methods. The availability as an user-friendly R package will also permit its routine application in clinical diagnostic and research laboratories.

## Conflict of interest

None declared. The views expressed are those of the authors and not necessarily those of the NHS or Department of Health.

## Acknowledgements

We are grateful to Petros Dellaportas for helpful discussions about INLA.

