## Supplementary Material for "MethylCal: Bayesian calibration of methylation levels"

### List of Figures

S.1 Schematic representation of MethylCal's models . . . S.2
S.2 Observed degree of bias introduced by PCR amplification of independent assays in two CGIs located on *SDHC* gene promoter (*SDHC* CpG:17 and *SDHC* CpG:27) . . . S.3
S.3 Observed degree of bias introduced by PCR amplification of independent assays in six imprinted DMRs (*PLAGL1*, *GRB10*, *MEST*, *H19*/*IGF2*, *KCNQ1OT1* and *MEG3*) . . . S.4
S.4 Methylation level of three independent assays in two CGIs located on *SDHC* gene promoter (*SDHC* CpG:17 and *SDHC* CpG:27) and one imprinted DMR (*PLAGL1*) calibrated by Moskalev's cubic polynomial regression S.5
S.5 Methylation level of independent assays in three imprinted DMRs (*GRB10*, *MEST* and *MEG3*) calibrated by Moskalev's cubic polynomial regression . . . S.6
S.6 Methylation level of three independent assays in two CGIs located on *SDHC* gene promoter (*SDHC* CpG:17 and *SDHC* CpG:27) and one imprinted DMR (*PLAGL1*) calibrated by MethylCal . . . S.7
S.7 Methylation level of independent assays in three imprinted DMRs (*GRB10*, *MEST* and *MEG3*) calibrated by MethylCal. . . . S.8
S.8 Posterior mean of MethylCal's random-effect AMP of eight independent assays in two CGIs located on *SDHC* gene promoter (*SDHC* CpG:17 and *SDHC* CpG:27) and six imprinted DMRs (*PLAGL1*, *GRB10*, *MEST*, *H19*/*IGF2*, *KCNQ1OT1* and *MEG3*) . . . S.9
S.9 Posterior mean of MethylCal's latent Gaussian field of two independent assays in one CGI located on *SDHC* gene promoter (*SDHC* CpG:17) and one imprinted DMR (*H19*/*IGF2*) . . . S.10
S.10 Posterior mean of MethylCal's random-intercepts CpG and random-slopes CpG$^*$ of six independent assays in one CGI located on *SDHC* gene promoter (*SDHC* CpG:27) and five imprinted DMRs (*PLAGL1*, *GRB10*, *MEST*, *KCNQ1OT1* and *MEG3*) . . . S.11
S.11 Corrected methylation degree of the *KCNQ1OT1* and *H19*/*IGF2* assays using Moskalev's cubic polynomial regression and MethylCal . . . S.12
S.12 Calibrated methylation level and corrected methylation degree of the *RELA* assay in celiac patients using Moskalev's cubic polynomial regression and MethylCal . . . S.13
S.13 Calibrated methylation level and corrected methylation degree of the *TNFAIP3* assay in celiac patients using Moskalev's cubic polynomial regression and MethylCal . . . S.14

### List of Tables

S.1 Goodness-of-fit performance of MethylCal and Moskalev's cubic polynomial regression on six independent assays in two CGIs located on *SDHC* gene promoter (*SDHC* CpG:17 and *SDHC* CpG:27) and four imprinted DMRs (*PLAGL1*, *GRB10*, *MEST* and *MEG3*) in the Beckwith-Wiedemann syndrome data set . . . S.15
S.2 Comparison of the goodness-of-fit performance between MethylCal's best model and Moskalev's cubic polynomial regression in the Beckwith-Wiedemann syndrome data set . . . S.16
S.3 Goodness-of-fit performance of MethylCal and Moskalev's cubic polynomial regression on independent assays in three genes (*NFKBIA*, *RELA*, and *TNFAIP3*) in the celiac data set . . . S.16

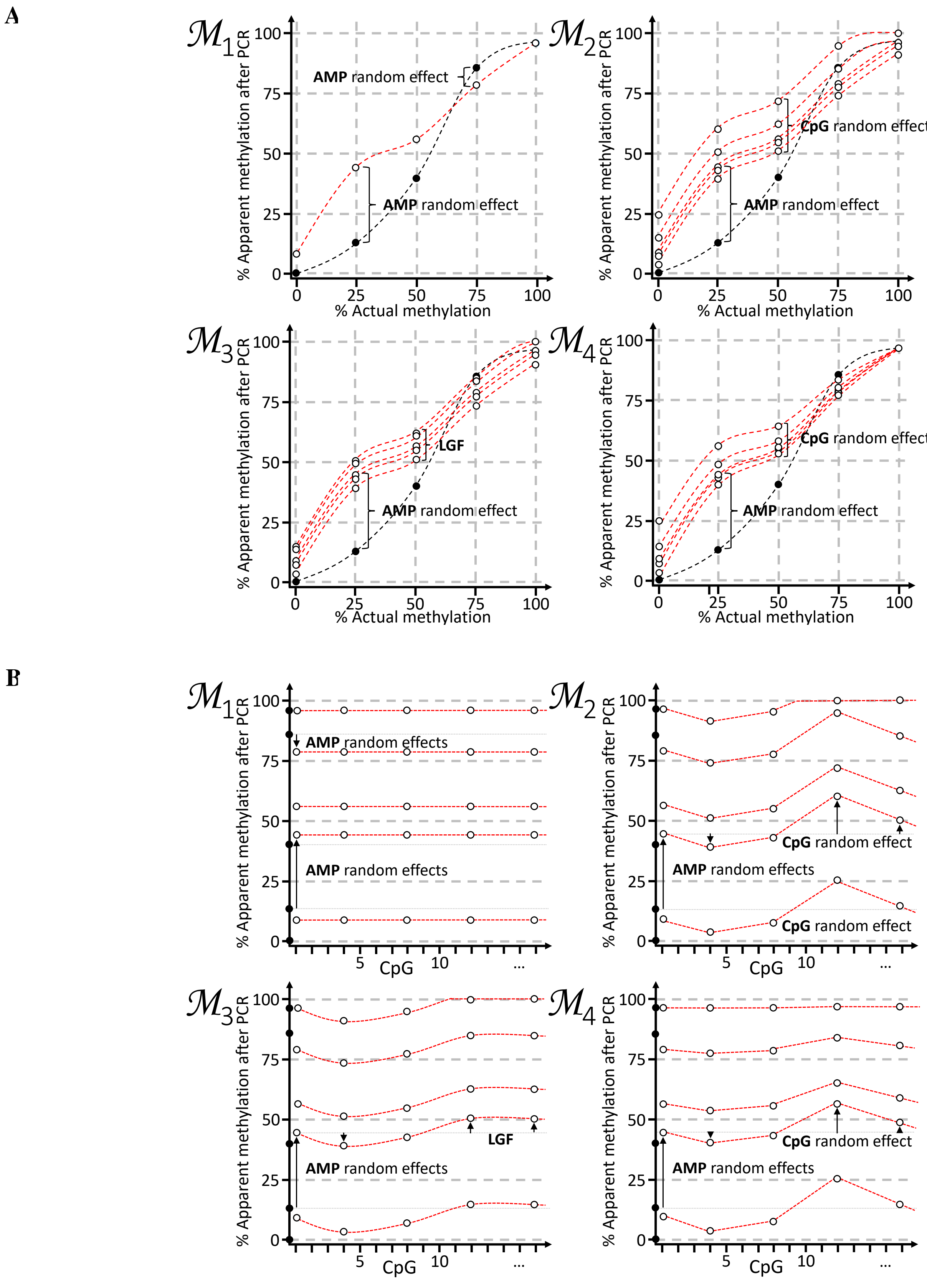


**Figure S.1.** Schematic representation of MethylCal's models described in the main text. (**A**) The apparent level of methylation observed after amplification ($y$-axis) is plotted as a function of the actual methylation percentages (AMP) ($x$-axis). The black dots and the black dotted line represent the apparent level of methylation predicted by the best-fit cubic polynomial regression (CPR) across all CpGs, whereas dots and the red dotted lines depict the level of methylation predicted by MethylCal's models. In model $\mathcal{M}_1$, the random-effect AMP is introduced and the level of methylation predicted by the CPR is shifted at each AMP by a different amount. In model $\mathcal{M}_2$, a random effect is added to model the variability of the apparent level of methylation after PCR across CpGs, with the level of methylation predicted by $\mathcal{M}_1$ shifted at each CpG independently. In model $\mathcal{M}_3$, the random-effect CpG is smoothed across CpGs by a latent Gaussian field (LGF). Finally, in model $\mathcal{M}_4$, the random-effects CpG depends on the AMP. (**B**) The apparent level of methylation observed after amplification ($y$-axis) is plotted as a function of the CpGs in the assay ($x$-axis). The black dots at $x = 0$ represent the level of methylation predicted by the best-fit CPR across all CpGs, whereas dots and the red dotted lines depict the level of methylation predicted by MethylCal's models.

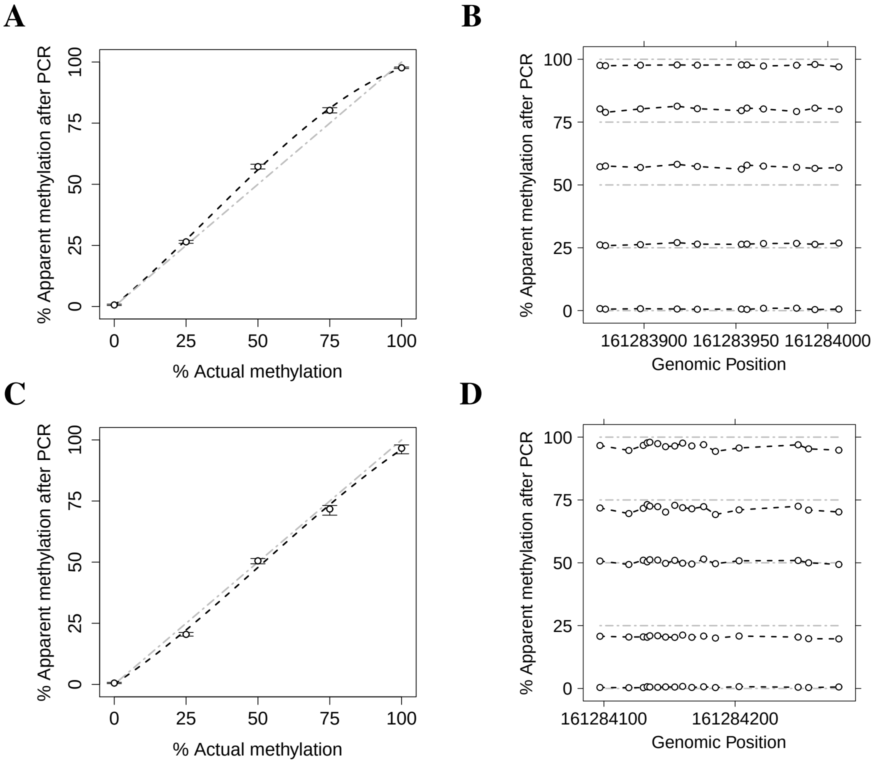


**Figure S.2.** Observed degree of bias introduced by PCR amplification of independent assays in two CGIs located on *SDHC* gene promoter ((**A**-**B**) *SDHC* CpG:17 and (**C**-**D**) *SDHC* CpG:27). (**A**-**C**) The apparent level of methylation observed after amplification ($y$-axis) is plotted as a function of the actual methylation percentage (AMP) ($x$-axis). For each AMP, the boxplot depicts the range and the median of the apparent level of methylation after PCR across CpGs. The black dashed line represents Moskalev's cubic polynomial interpolation curve of the average (across CpGs) apparent level of methylation after PCR. The grey dot-dashed line represents an unbiased plot. (**B**-**D**) The apparent level of methylation after PCR ($y$-axis) is plotted (circles) as a function of the CpGs genomic position in the assay ($x$-axis). The grey dot-dashed lines depict unbiased apparent levels of methylation after PCR.

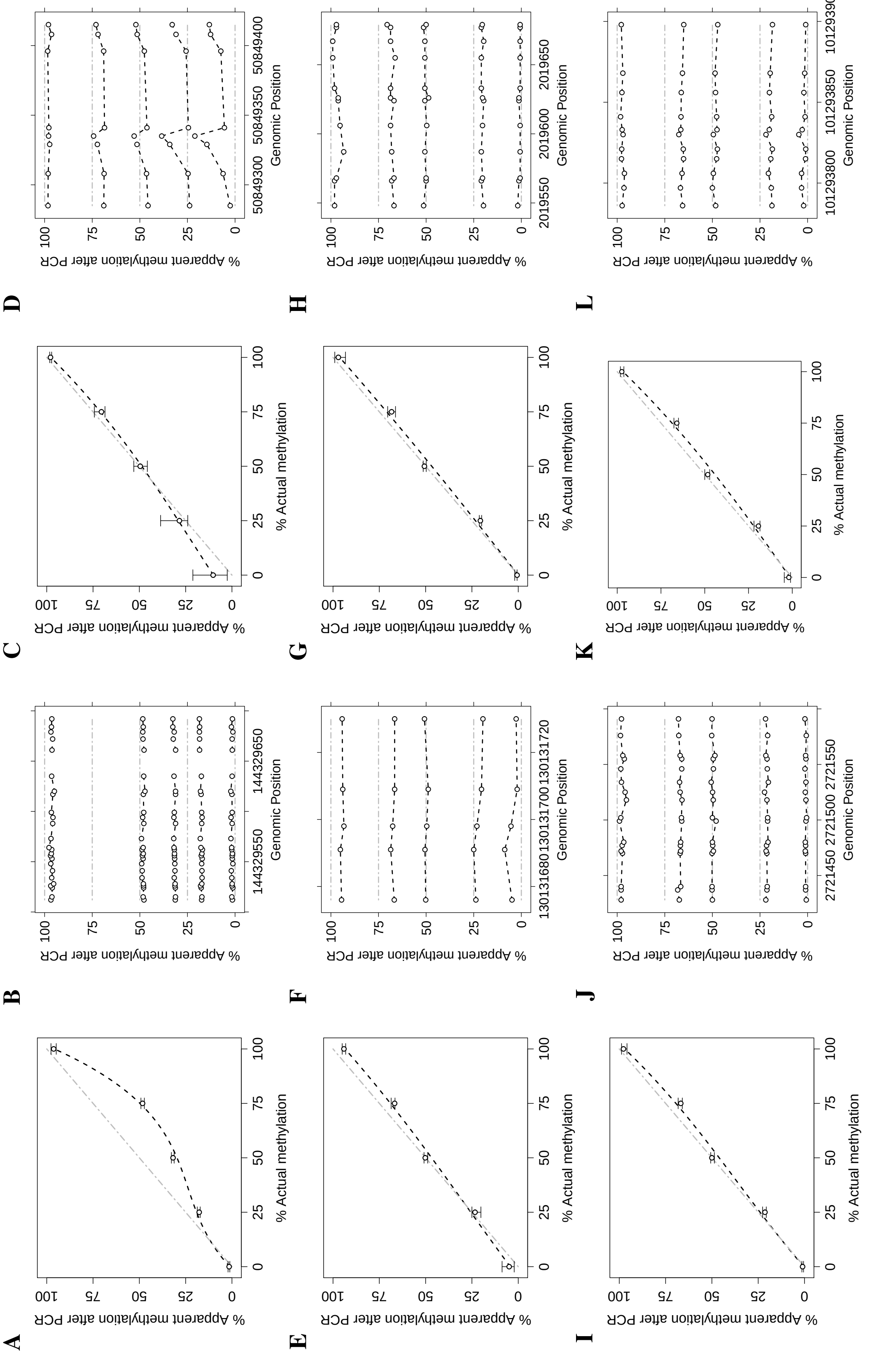


**Figure S.3.** Observed degree of bias introduced by PCR amplification of independent assays in six imprinted DMRs ((**A**-**B**) *PLAGL1*, (**C**-**D**) *GRB10*, (**E**-**F**) *MEST*, (**G**-**H**) *H19*/*IGF2*, (**I**-**J**) *KCNQ1OT1* and (**K**-**L**) *MEG3*). (**A**-**C**-**E**-**G**-**I**-**K**) The apparent level of methylation observed after amplification ($y$-axis) is plotted as a function of the actual methylation percentage (AMP) ($x$-axis). For each AMP, the boxplot depicts the range and the median of the apparent level of methylation after PCR across CpGs. The black dashed line represents Moskalev's cubic polynomial interpolation curve of the average (across CpGs) apparent level of methylation after PCR. The grey dot-dashed line represents an unbiased plot. (**B**-**D**-**F**-**H**-**J**-**L**) The apparent level of methylation after PCR ($y$-axis) is plotted (circles) as a function of the CpGs genomic position in the assay ($x$-axis). The grey dot-dashed lines depict unbiased apparent levels of methylation after PCR. In the *PLAGL1* assay (**A**-**B**) three CpGs were removed after visual inspection since their apparent level of methylation observed after amplification were very different from the other CpGs in the assay.

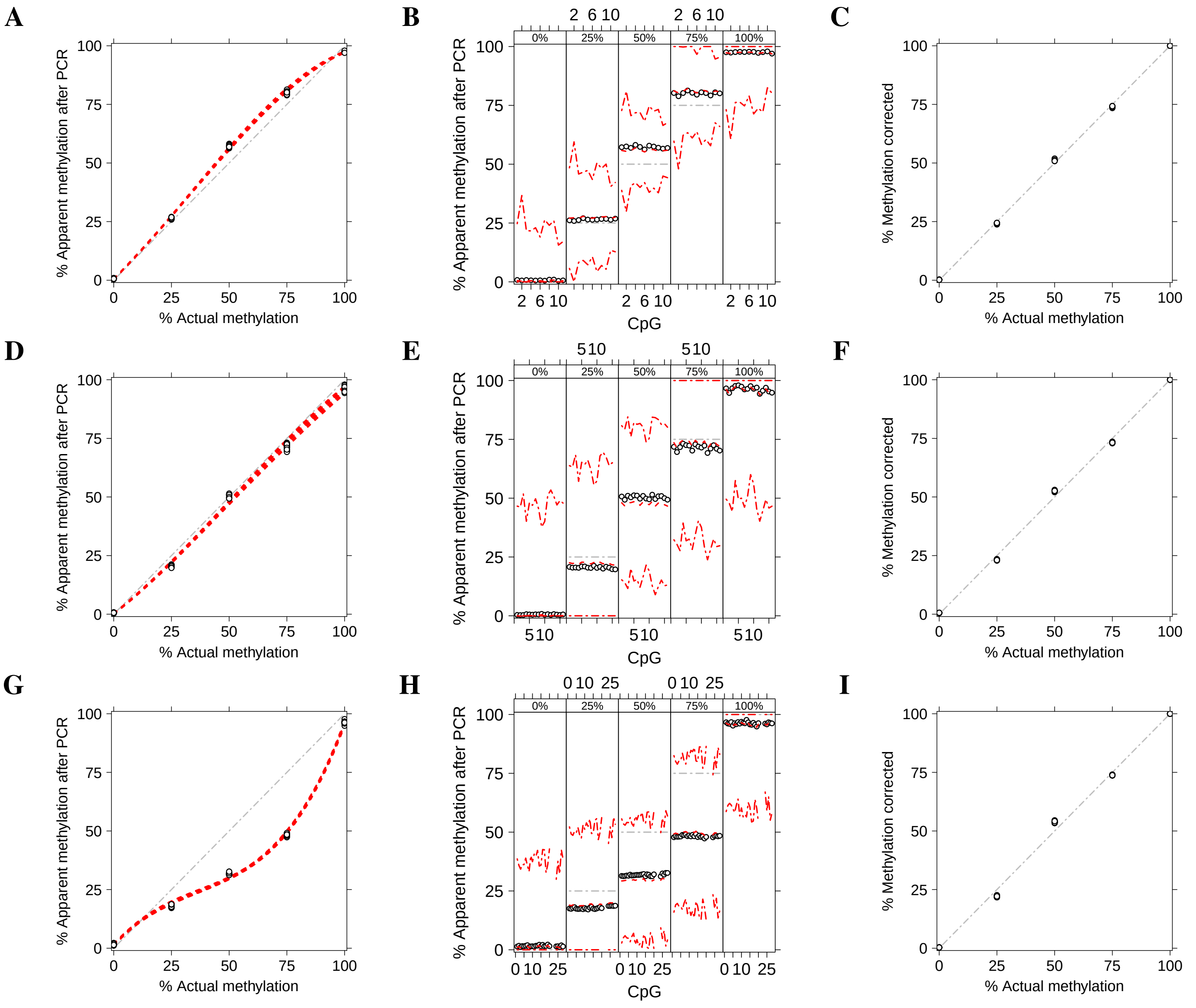


**Figure S.4.** Methylation level of three independent assays in two CGIs located on *SDHC* gene promoter ((**A**-**B**-**C**) *SDHC* CpG:17 and (**D**-**E**-**F**) *SDHC* CpG:27) and one imprinted DMR ((**G**-**H**-**I**) *PLAGL1*) calibrated by Moskalev's cubic polynomial regression (CPR). (**A**-**D**-**G**) The apparent level of methylation observed after amplification ($y$-axis) is plotted as a function of the actual methylation percentage (AMP) ($x$-axis). Circles depict the apparent level of methylation for each CpG at different AMP, whereas the red dotted lines show Moskalev's CPR for each CpG. The grey dash-dotted line represents an unbiased plot. (**B**-**E**-**H**) The apparent level of methylation observed after amplification ($y$-axis) is plotted (circles) as a function of the CpGs in the assay ($x$-axis), stratified by AMP (top figure box and the grey dot-dashed line). For each stratum, the red dotted line shows the level of methylation predicted by Moskalev's CPR, whereas the red dot-dashed lines depict the 95% predicted confidence interval. (**C**-**F**-**I**) The result of PCR-bias correction by using Moskalev's CPR. The corrected methylation degree ($y$-axis) is plotted as a function of the AMP ($x$-axis). The grey dash-dotted line represents a perfectly corrected plot.

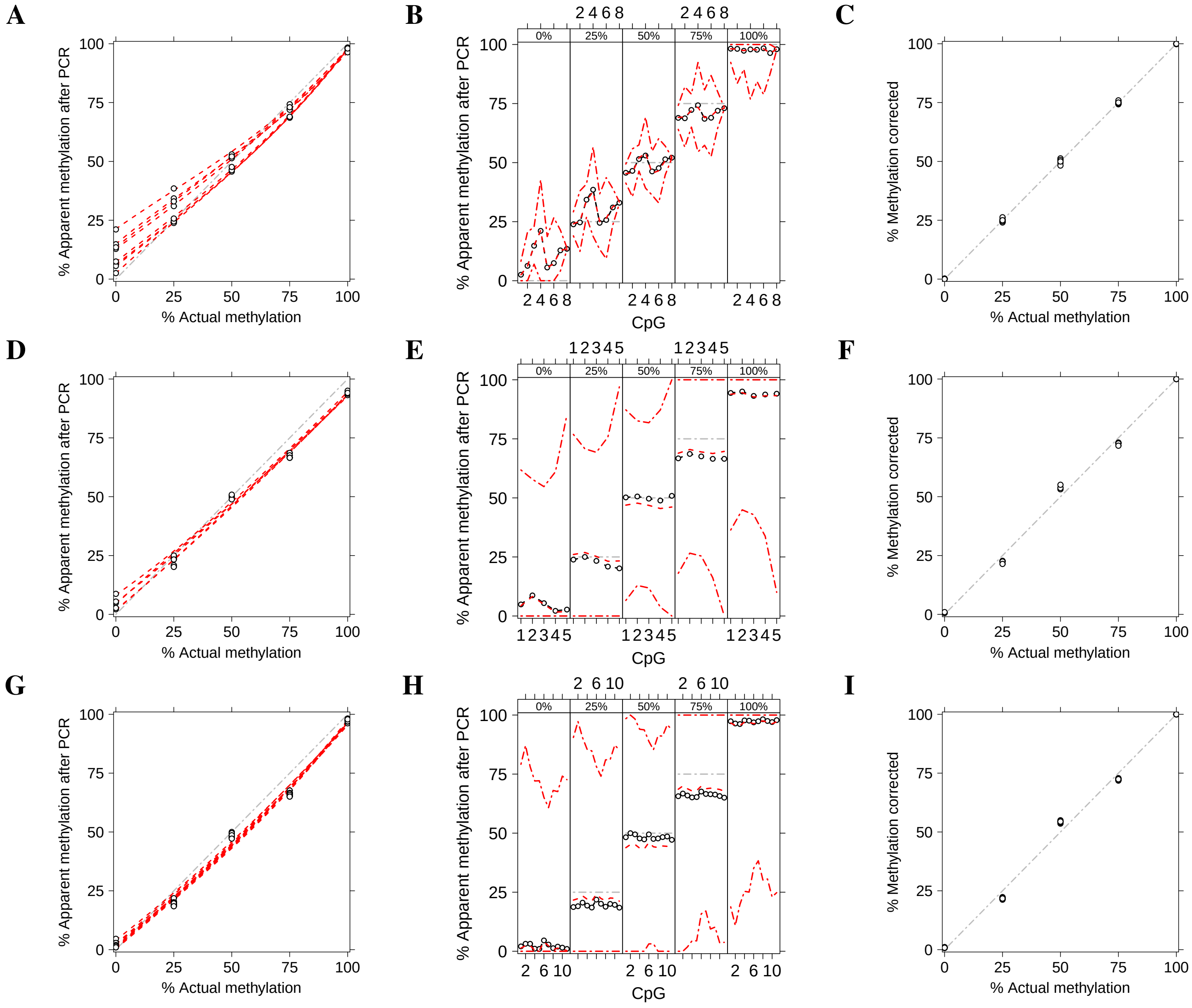


**Figure S.5.** Methylation level of independent assays in three imprinted DMRs ((**A**-**B**-**C**) *GRB10*, (**D**-**E**-**F**) *MEST* and (**G**-**H**-**I**) *MEG3*) calibrated by Moskalev's cubic polynomial regression (CPR). (**A**-**D**-**G**) The apparent level of methylation observed after amplification ($y$-axis) is plotted as a function of the actual methylation percentage (AMP) ($x$-axis). Circles depict the apparent level of methylation for each CpG at different AMP, whereas the red dotted lines show Moskalev's CPR for each CpG. The grey dash-dotted line represents an unbiased plot. (**B**-**E**-**H**) The apparent level of methylation observed after amplification ($y$-axis) is plotted (circles) as a function of the CpGs in the assay ($x$-axis), stratified by AMP (top figure box and the grey dot-dashed line). For each stratum, the red dotted line shows the level of methylation predicted by Moskalev's CPR, whereas the red dot-dashed lines depict the 95% predicted confidence interval. (**C**-**F**-**I**) The result of PCR-bias correction by using Moskalev's CPR. The corrected methylation degree ($y$-axis) is plotted as a function of the AMP ($x$-axis). The grey dash-dotted line represents a perfectly corrected plot.

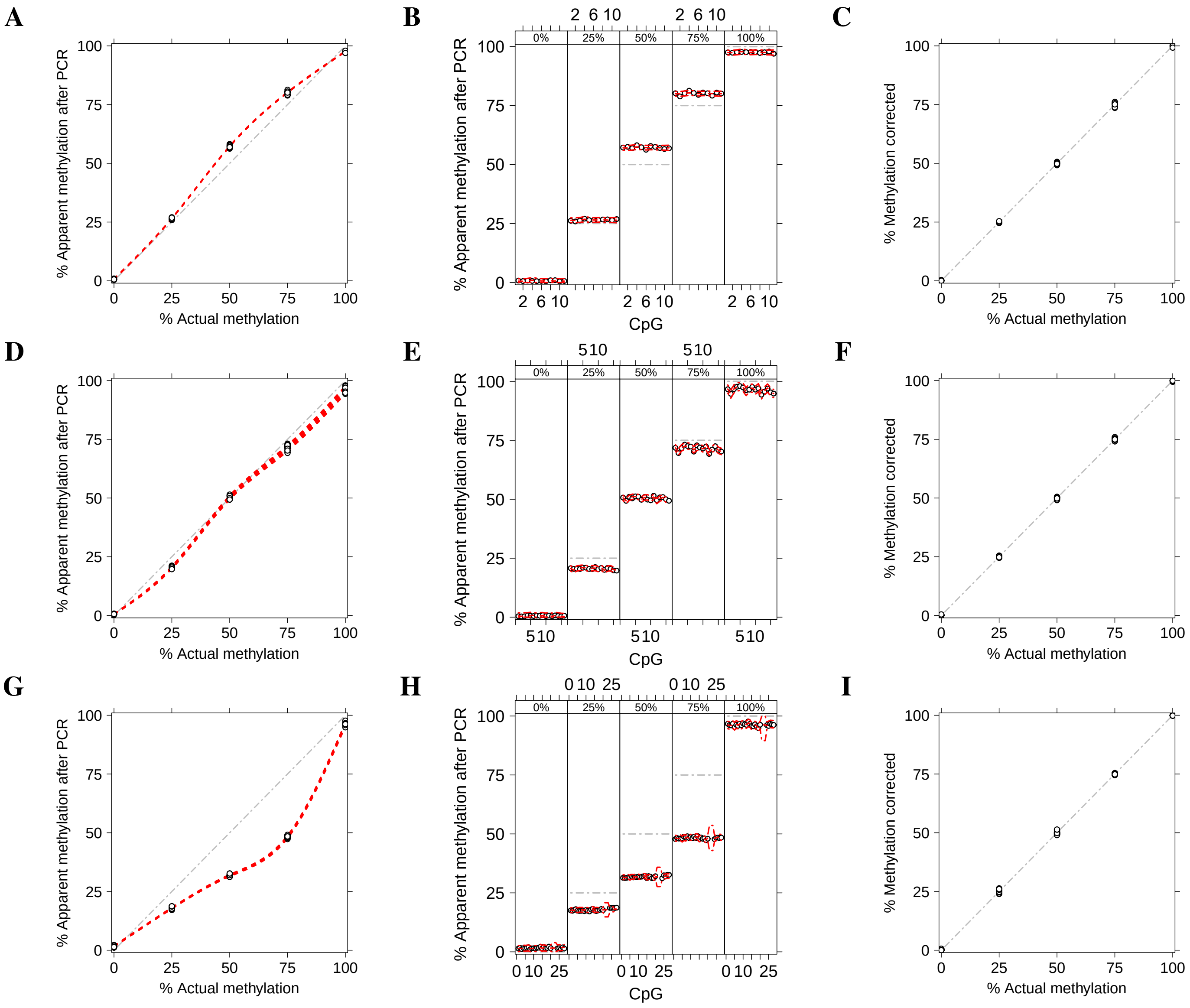


**Figure S.6.** Methylation level of three independent assays in two CGIs located on *SDHC* gene promoter ((**A**-**B**-**C**) *SDHC* CpG:17 and (**D**-**E**-**F**) *SDHC* CpG:27) and one imprinted DMR ((**G**-**H**-**I**) *PLAGL1*) calibrated by MethylCal. In the assay *SDHC* CpG:17, model $\mathcal{M}_3$ is selected by the DIC, whereas model $\mathcal{M}_4$ is selected in the assays *SDHC* CpG:27 and *PLAGL1*. (**A**-**D**-**G**) The apparent level of methylation observed after amplification ($y$-axis) is plotted as a function of the actual methylation percentage (AMP) ($x$-axis). Circles depict the apparent level of methylation for each CpG at different AMPs, whereas the red dotted lines show MethylCal's predicted methylation for each CpG. The grey dash-dotted line represents an unbiased plot. (**B**-**E**-**H**) The apparent level of methylation observed after amplification ($y$-axis) is plotted (circles) as a function of the CpGs in the assay ($x$-axis), stratified by AMP (top figure box and the grey dot-dashed line). For each stratum, the red dotted line shows the level of methylation predicted by MethylCal, whereas the red dot-dashed lines depict the 95% predicted credible interval. (**C**-**F**-**I**) The result of PCR-bias correction obtained by using MethylCal. The corrected methylation degree ($y$-axis) is plotted as a function of the AMP ($x$-axis). The grey dash-dotted line represents a perfectly corrected plot.

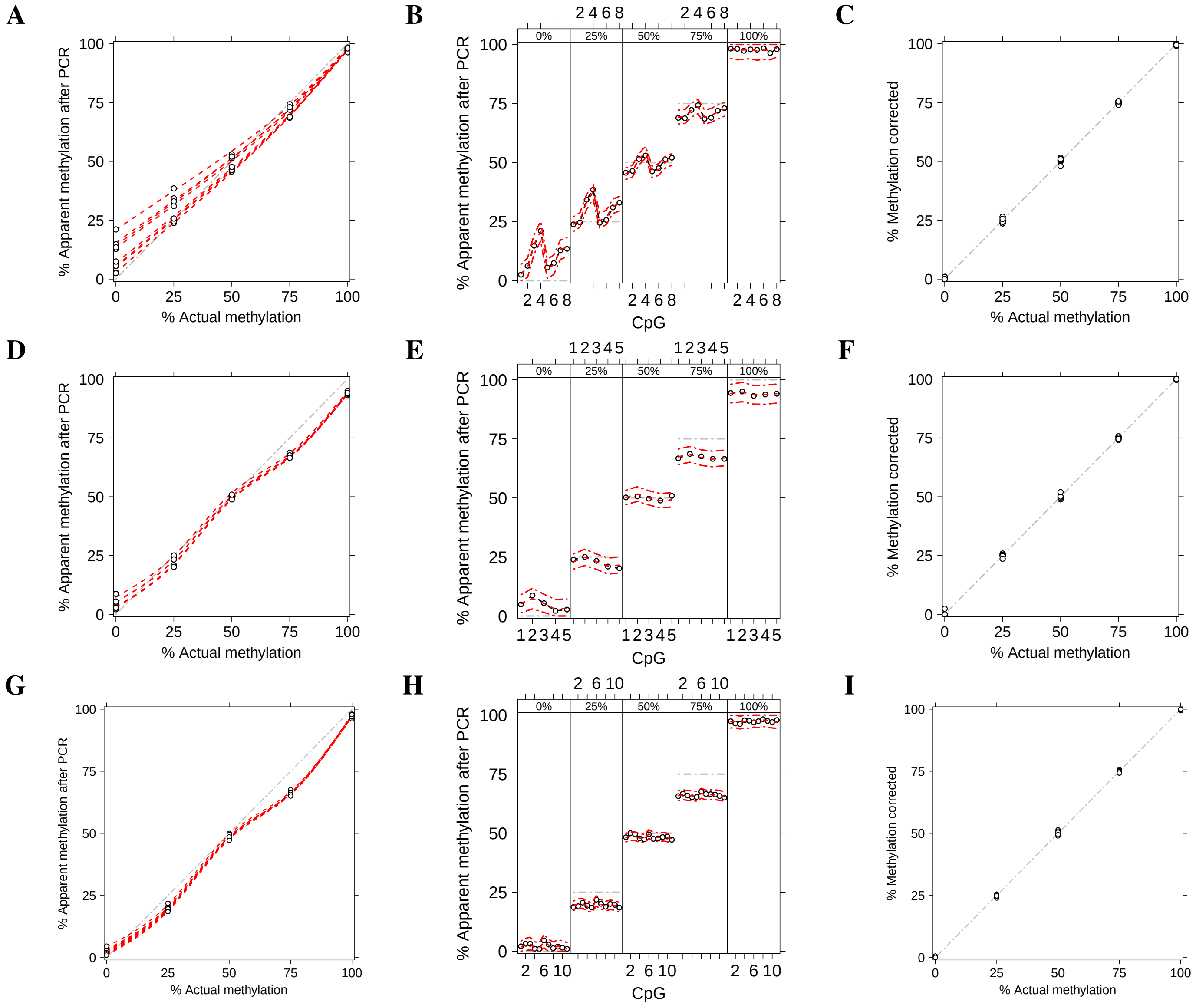


**Figure S.7.** Methylation level of independent assays in three imprinted DMRs ((**A**-**B**-**C**) *GRB10*, (**D**-**E**-**F**) *MEST* and (**G**-**H**-**I**) *MEG3*) calibrated by MethylCal. In all assays analysed, model $\mathcal{M}_4$ is selected by the DIC. (**A**-**D**-**G**) The apparent level of methylation observed after amplification ($y$-axis) is plotted as a function of the actual methylation percentage (AMP) ($x$-axis). Circles depict the apparent level of methylation for each CpG at different AMPs, whereas the red dotted lines show MethylCal's predicted methylation for each CpG. The grey dash-dotted line represents an unbiased plot. (**B**-**E**-**H**) The apparent level of methylation observed after amplification ($y$-axis) is plotted (circles) as a function of the CpGs in the assay ($x$-axis), stratified by AMP (top figure box and the grey dot-dashed line). For each stratum, the red dotted line shows the level of methylation predicted by MethylCal, whereas the red dot-dashed lines depict the 95% predicted credible interval. (**C**-**F**-**I**) The result of PCR-bias correction obtained by using MethylCal. The corrected methylation degree ($y$-axis) is plotted as a function of the AMP ($x$-axis). The grey dash-dotted line represents a perfectly corrected plot.

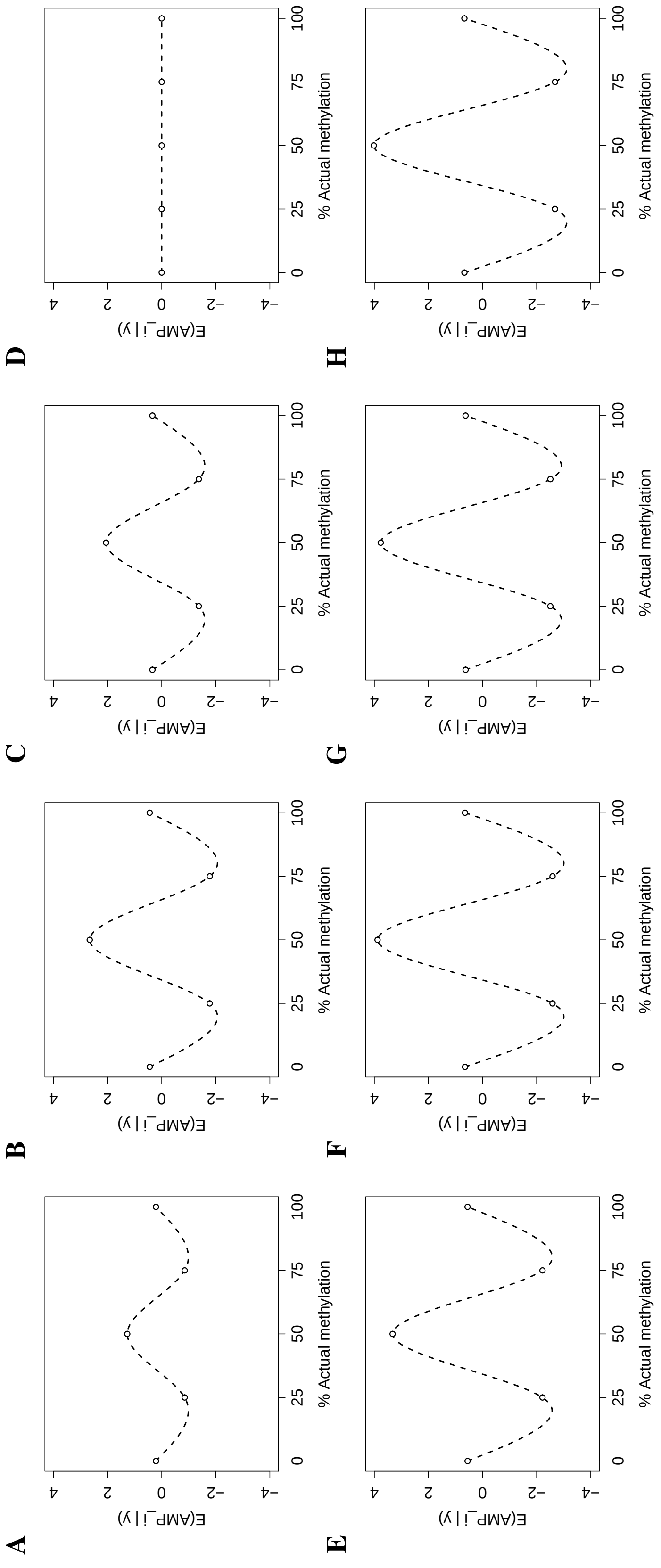


**Figure S.8.** Posterior mean of MethylCal's random-effect AMP (circles) and cubic spline interpolation (dashed black line) of eight independent assays in two CGIs located on *SDHC* gene promoter ((**A**) *SDHC* CpG:17 and (**B**) *SDHC* CpG:27) and six imprinted DMRs ((**C**) *PLAGL1*, (**D**) *GRB10* (**E**) *MEST*, (**F**) *H19*/*IGF2*, (**G**) *KCNQ1OT1* and (**H**) *MEG3*). In assay *GRB10*, no AMP random effects were detected.

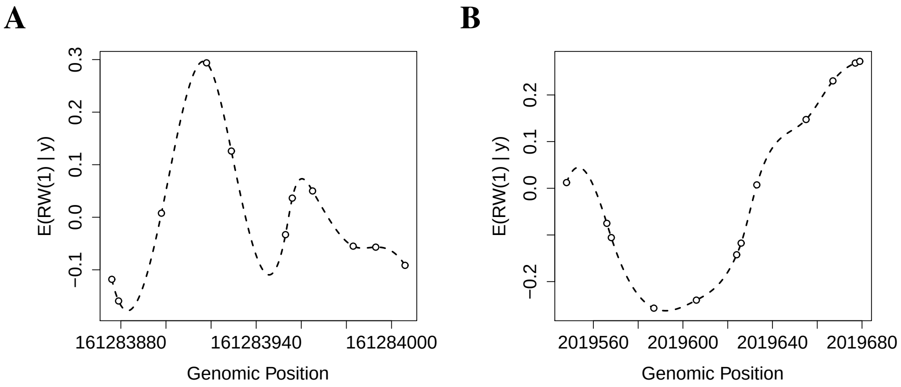


**Figure S.9.** Posterior mean of MethylCal's latent Gaussian field (LGF) at observed CpGs (dots) and LGF interpolation (dashed black line) of two independent assays in one CGI located on *SDHC* gene promoter ((**A**) *SDHC* CpG:17) and one imprinted DMR ((**B**) *H19*/*IGF2*).

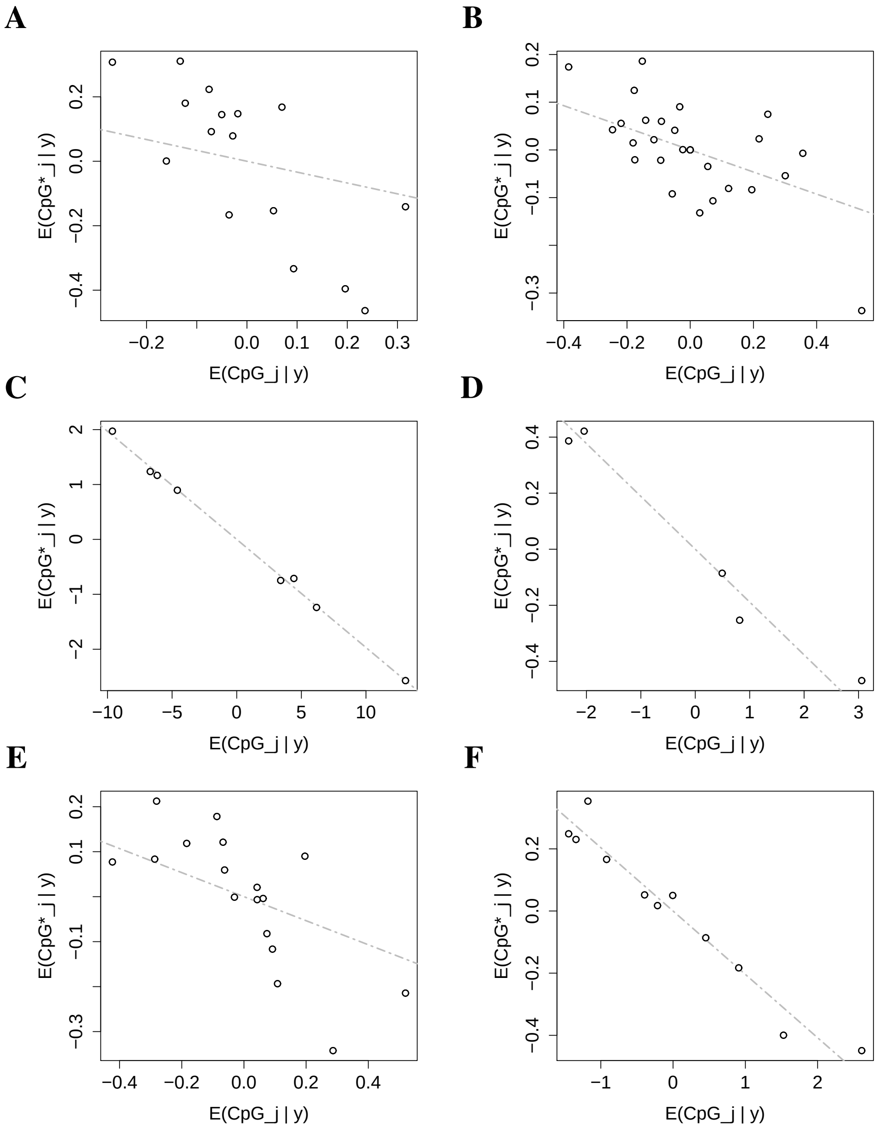


**Figure S.10.** Scatterplot of the posterior mean of MethylCal's random-intercepts $\text{CpG}_j$ and random-slopes $\text{CpG}_j^*$ (circles) of six independent assay in one CGI located on *SDHC* gene promoter ((**A**) SDHC CpG:27) and five imprinted DMRs ((**B**) *PLAGL1*, (**C**) *GRB10*, (**D**) *MEST*, (**E**) *KCNQ1OT1* and (**F**) *MEG3*). The grey dash-dotted line depicts the posterior mean of the covariance $\Sigma_{12}$ between the random-intercepts $\text{CpG}_j$ and random-slopes $\text{CpG}_j^*$ (see main text for details). $\mathrm{E}(\Sigma_{12}|\boldsymbol{y})$ can be also interpreted as the "robustified" covariance estimate between the two random effects, see panels (**A**), (**B**) and (**E**).

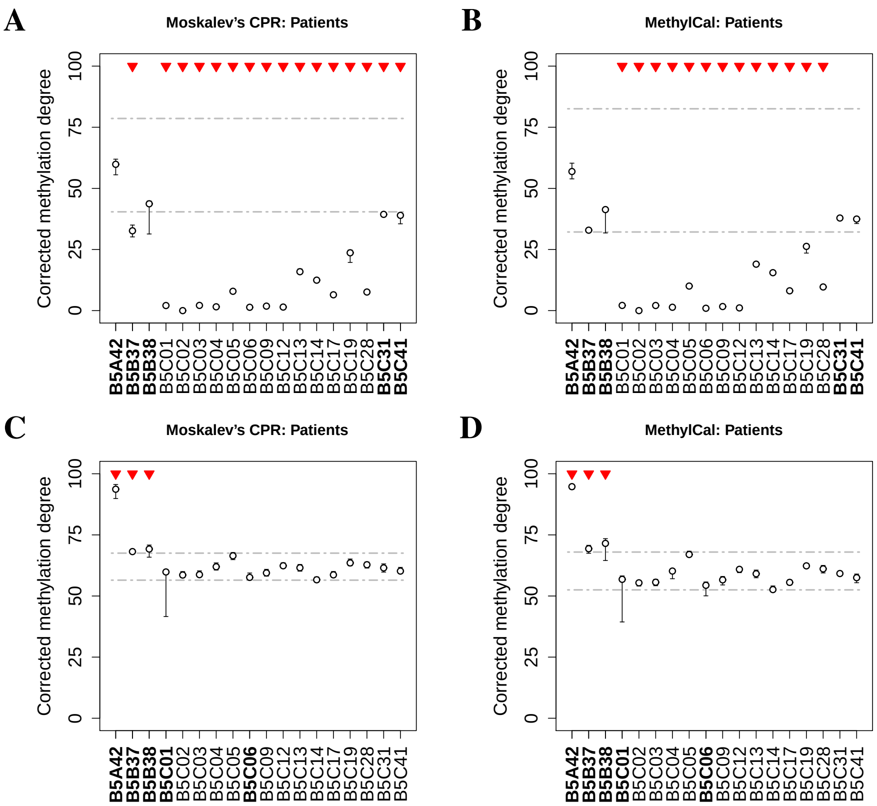


**Figure S.11.** Corrected methylation degree of the (**A**-**B**) *KCNQ1OT1* and (**C**-**D**) *H19*/*IGF2* assays using Moskalev's cubic polynomial regression (left panels) and MethylCal (right panels) for patients potentially affected by Beckwith-Wiedemann syndrome. For each patient ($x$-axis), the boxplot depicts the range and the median of the corrected methylation degree ($y$-axis) across CpGs. The dashed-dotted grey lines show the $\alpha = 0.01$ confidence interval centered around the overall mean (see main text for details). Top red triangles indicate patients classified as having undergone loss or gain of methylation if their average (across CpGs) corrected methylation degree is outside the healthy controls' confidence interval. Bold font ($x$-axis) indicates patients' classification described in the main text.

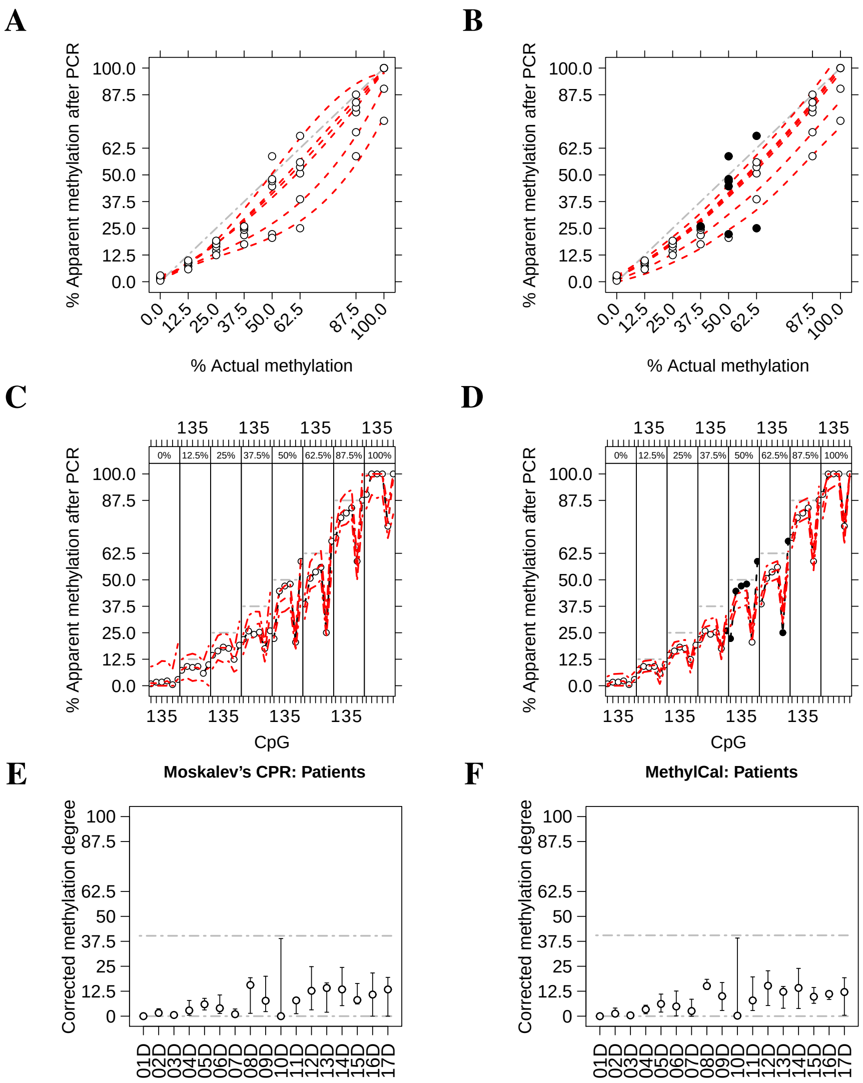


**Figure S.12.** Calibrated methylation level and corrected methylation degree of the *RELA* assay in celiac patients using Moskalev's cubic polynomial regression (left panels) and MethylCal (right panels). (**A**-**B**) The apparent level of methylation observed after amplification ($y$-axis) is plotted as a function of the actual methylation percentage (AMP) ($x$-axis). Circles depict the apparent level of methylation after PCR for each CpG at different AMPs, whereas the red dotted lines show MethylCal's predicted level of methylation for each CpG. Black dots highlight potential outliers. The grey dash-dotted line represents an unbiased plot. (**C**-**D**) The apparent level of methylation observed after amplification ($y$-axis) is plotted (circles) as a function of the CpGs in the DMR ($x$-axis), stratified by AMP (top figure box and the grey dot-dashed line). For each stratum, the red dotted line shows the level of methylation predicted by MethylCal, whereas the red dot-dashed lines depict the 95% prediction credible interval. (**E**-**F**) For each patient ($x$-axis), the boxplot depicts the range and the mean (circle) of the corrected methylation degree ($y$-axis) across CpGs while the dashed-dotted grey lines show the healthy controls' confidence interval.

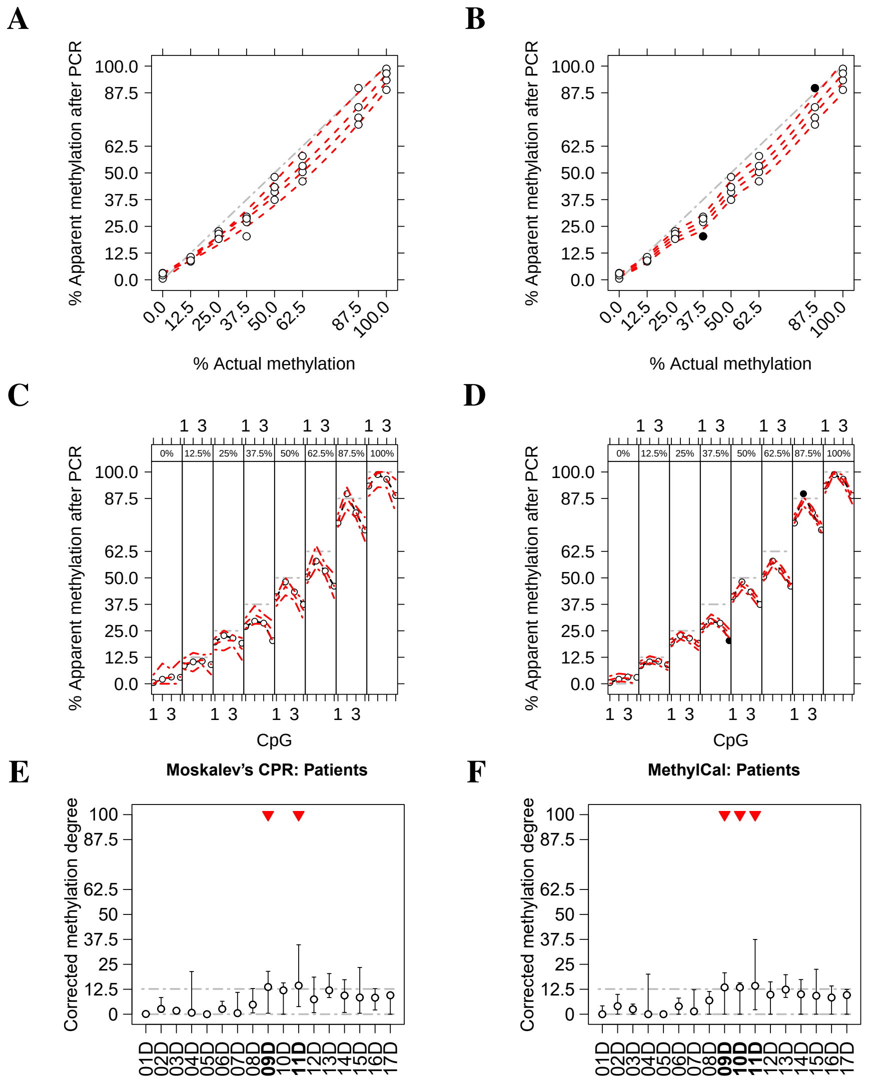


**Figure S.13.** Calibrated methylation level and corrected methylation degree of the *TNFAIP3* assay in celiac patients using Moskalev's cubic polynomial regression (left panels) and MethylCal (right panels). (**A**-**B**) The apparent level of methylation observed after amplification ($y$-axis) is plotted as a function of the actual methylation percentage (AMP) ($x$-axis). Circles depict the apparent level of methylation after PCR for each CpG at different AMPs, whereas the red dotted lines show MethylCal's predicted level of methylation for each CpG. Black dots highlight potential outliers. The grey dash-dotted line represents an unbiased plot. (**C**-**D**) The apparent level of methylation observed after amplification ($y$-axis) is plotted (circles) as a function of the CpGs in the DMR ($x$-axis), stratified by AMP (top figure box and the grey dot-dashed line). For each stratum, the red dotted line shows the level of methylation predicted by MethylCal, whereas the red dot-dashed lines depict the 95% prediction credible interval. (**E**-**F**) For each patient ($x$-axis), the boxplot depicts the range and the mean (circle) of the corrected methylation degree ($y$-axis) across CpGs while the dashed-dotted grey lines show the healthy controls' confidence interval. Top red triangles indicate patients classified as having undergone loss or gain of methylation if their average (across CpGs) corrected methylation degree is outside the healthy controls' confidence interval.

| | | | Moskalev | MethylCal | | | |
|---|---|---|---|---|---|---|---|
| | | | | $\mathcal{M}_1$ | $\mathcal{M}_2$ | $\mathcal{M}_3$ | $\mathcal{M}_4$ |
| | | DIC | – | 74.00 | 73.65 | **71.41** | 75.30 |
| | | RSS | 37.99 | 9.88 | 9.87 | 7.90 | **5.68** |
| *SDHC* CpG:17 | CpGs & AMPs | MSEP | 55.83 | | | **0.21** | |
| | | CV-index | -256.10 | | | **0.03** | |
| | CpGs | MSEP | 4.66 | | | **1.15** | |
| | | CV-index | -3.29 | | | **-0.06** | |
| | AMPs | MSEP | 614.16 | | | **583.25** | |
| | | DIC | – | 203.29 | 163.90 | 162.83 | **109.35** |
| | | RSS | 223.03 | 50.98 | 19.82 | 22.84 | **8.78** |
| *SDHC* CpG:27 | CpGs & AMPs | MSEP | 225.48 | | | | **0.25** |
| | | CV-index | -310.09 | | | | **0.65** |
| | CpGs | MSEP | 18.74 | | | | **3.62** |
| | | CV-index | -4.17 | | | | **0.00** |
| | AMPs | MSEP | 3,607.74 | | | | **3,580.19** |
| | | DIC | – | 167.06 | 167.10 | 154.13 | **151.39** |
| | | RSS | 208.62 | 25.27 | 25.23 | 20.11 | **12.17** |
| *PLAGL1* | CpGs & AMPs | MSEP | 120.72 | | | | **0.19** |
| | | CV-index | -615.26 | | | | **0.02** |
| | CpGs | MSEP | 8.53 | | | | **0.98** |
| | | CV-index | -7.71 | | | | **0.00** |
| | AMPs | MSEP | 3,380.11 | | | | **3,354.38** |
| | | DIC | – | 230.48 | 202.71 | 206.13 | **122.55** |
| | | RSS | **8.73** | 576.72 | 194.53 | 217.55 | 18.40 |
| *GRB10* | CpGs & AMPs | MSEP | 22.04 | | | | **1.57** |
| | | CV-index | -0.17 | | | | **0.92** |
| | CpGs | MSEP | **53.26** | | | | 93.80 |
| | | CV-index | **0.43** | | | | 0.00 |
| | AMPs | MSEP | 176.33 | | | | **102.85** |
| | | DIC | – | 102.22 | 101.64 | 91.48 | **78.75** |
| | | RSS | 117.68 | 52.01 | 25.43 | 26.57 | **9.99** |
| *MEST* | CpGs & AMPs | MSEP | 385.26 | | | | **1.53** |
| | | CV-index | -119.10 | | | | **0.52** |
| | CpGs | MSEP | 32.72 | | | | **16.01** |
| | | CV-index | -1.04 | | | | **0.00** |
| | AMPs | MSEP | 1,926.28 | | | | **1,848.71** |
| | | DIC | – | 155.11 | 147.41 | 149.14 | **125.45** |
| | | RSS | 351.82 | 43.21 | 29.86 | 32.92 | **12.18** |
| *MEG3* | CpGs & AMPs | MSEP | 517.26 | | | | **0.67** |
| | | CV-index | -543.40 | | | | **0.29** |
| | CpGs | MSEP | 38.33 | | | | **4.75** |
| | | CV-index | -7.07 | | | | **0.00** |
| | AMPs | MSEP | 5,689.90 | | | | **5,639.63** |

**Table S.1.** Goodness-of-fit performance of MethylCal and Moskalev's cubic polynomial regression (CPR) on six independent assays in two CGIs located on *SDHC* gene promoter (*SDHC* CpG:17 and *SDHC* CpG:27) and four imprinted DMRs (*PLAGL1*, *GRB10*, *MEST* and *MEG3*) in the Beckwith-Wiedemann syndrome data set. For each MethylCal's model the Deviance Information Criterion (DIC) is calculated and for Moskalev's CPR also the Residual Sum of Squares (RSS). The Mean Squared Error of Prediction (MSEP) and the CV-index are evaluated for the best MethylCal's model (based on the DIC) and for Moskalev's CPR when the leave-one-out cross-validation is performed across (i) CpGs *and* actual methylation percentages (AMPs); (ii) only CpGs; (iii) only AMPs. For each predictive measure, the best result is highlighted in bold.

| | | MethylCal *vs* Moskalev | MethylCal's CV-index $\geq 0$ |
|---|---|---|---|
| | RSS | 7/8 | - |
| CpGs & AMPs | MSEP | 8/8 | - |
| | CV-index | 8/8 | 7/8 |
| CpGs | MSEP | 7/8 | - |
| | CV-index | 7/8 | 7/8 |
| AMPs | MSEP | 8/8 | - |

**Table S.2.** Comparison of the "in-sample" and "out-of-sample" goodness-of-fit performance between MethylCal's best model selected by the DIC and Moskalev's cubic polynomial regression (CPR) across the assays considered in the Beckwith-Wiedemann syndrome data set. For the two methods considered, the Residual Sum of Squares (RSS), the Mean Squared Error of Prediction (MSEP) and the CV-index are calculated with the leave-one-out cross-validation performed across: (i) CpGs *and* actual methylation percentages (AMPs); (ii) CpGs; (iii) AMPs. The number of times MethylCal's best model performs better than or equal to Moskalev's CPR is reported as well as the number of times it is non-negative.

| | | | Moskalev | MethylCal | | | |
|---|---|---|---|---|---|---|---|
| | | | | $\mathcal{M}_1$ | $\mathcal{M}_2$ | $\mathcal{M}_3$ | $\mathcal{M}_4$ |
| *NFKBIA* | | DIC | - | 307.5 | 273.25 | 274.59 | 249.12 |
| | | RSS | **241.66** | 3,929.12 | 1,163.32 | 1,211.34 | 658.08 (83.93) |
| | CpGs & AMPs | MSEP | 31.00 | | | | **29.86** |
| | | CV-index | 0.79 | | | | **0.80** |
| | CpGs | MSEP | **1,010.50** | | | | 1,151.91 |
| | | CV-index | **0.15** | | | | 0.03 |
| | AMPs | MSEP | 154.99 | | | | **135.09** |
| *RELA* | | DIC | | 353.85 | 322.51 | 324.11 | **291.33** |
| | | RSS | **392.61** | 3,583.22 | 1,417.73 | 1,472.86 | 724.92 (95.28) |
| | CpGs & AMPs | TNFAIP3 | 33.72 | | | | **23.80** |
| | | CV-index | 0.68 | | | | **0.77** |
| | CpGs | MSEP | 150.46 | | | | **92.86** |
| | | CV-index | -0.27 | | | | **0.22** |
| | AMPs | MSEP | 202.32 | | | | **162.57** |
| *TNFAIP3* | | DIC | | 188.34 | 167.84 | 168.39 | **136.59** |
| | | RSS | 83.81 | 478.66 | 200.54 | 201.76 | **53.75** (21.16) |
| | CpGs & AMPs | MSEP | 7.38 | | | | **4.12** |
| | | CV-index | 0.69 | | | | **0.82** |
| | CpGs | MSEP | 221.13 | | | | **197.83** |
| | | CV-index | -0.18 | | | | **-0.05** |
| | AMPs | MSEP | 29.51 | | | | **27.06** |

**Table S.3.** Goodness-of-fit performance of MethylCal and Moskalev's cubic polynomial regression (CPR) on independent assays in three genes (*NFKBIA*, *RELA*, and *TNFAIP3*) in the celiac data set. For each MethylCal's model the Deviance Information Criterion (DIC) is calculated and for MethylCal's models and Moskalev's CPR also the Residual Sum of Squares (RSS) (in parenthesis MethylCal's RSS after removing outliers). The Mean Squared Error of Prediction (MSEP) and the CV-index are evaluated for the best MethylCal's model (based on the DIC) and for Moskalev's CPR when the leave-one-out cross-validation is performed across (i) CpGs *and* actual methylation percentages (AMPs); (ii) only CpGs; (iii) only AMPs. For each predictive measure, the best result is highlighted in bold.
